# A modular approach for modeling the cell cycle based on functional response curves

**DOI:** 10.1101/2021.04.28.441747

**Authors:** Jolan De Boeck, Jan Rombouts, Lendert Gelens

## Abstract

Modeling biochemical reactions by means of differential equations often results in systems with a large number of variables and parameters. As this might complicate the interpretation and generalization of the obtained results, it is often desirable to reduce the complexity of the model. One way to accomplish this is by replacing the detailed reaction mechanisms of certain modules in the model by a mathematical expression that qualitatively describes the dynamical behavior of these modules. Such an approach has been widely adopted for ultrasensitive responses, for which underlying reaction mechanisms are often replaced by a single Hill equation. Also time delays are usually accounted for by using an explicit delay in delay differential equations. In contrast, however, S-shaped response curves, which are often encountered in bistable systems, are not easily modeled in such an explicit way. Here, we extend the classical Hill function into a mathematical expression that can be used to describe both ultrasensitive and S-shaped responses. We show how three ubiquitous modules (ultrasensitive responses, S-shaped responses and time delays) can be combined in different configurations and explore the dynamics of these systems. As an example, we apply our strategy to set up a model of the cell cycle consisting of multiple bistable switches, which can account for events such as DNA damage and coupling to the circadian clock in a phenomenological way.

**Author summary:** Bistability plays an important role in many biochemical processes and typically emerges from complex interaction patterns such as positive and double negative feedback loops. Here, we propose to theoretically study the effect of bistability in a larger interaction network. We explicitly incorporate a functional expression describing an S-shaped input-output curve in the model equations, without the need for considering the underlying biochemical events. This expression can be converted into a functional module for an ultrasensitive response, and a time delay is easily included as well. Exploiting the fact that several of these modules can easily be combined in larger networks, we construct a cell cycle model consisting of multiple bistable switches and show how this approach can account for a number of known properties of the cell cycle.

## Introduction

Cell division and the correct separation of genomic material among daughter cells is fundamental for the proper development, growth and reproduction of a living organism. The molecular mechanisms that underlie these processes are highly evolutionarily conserved. Incorrect cell division can have detrimental effects, ranging from developmental defects to the transformation of healthy somatic cells into cancer cells. Because of this, tight regulatory mechanisms are established early in embryonic development to ensure the correct replication of DNA and cell division. These control mechanisms, or cellular checkpoints, ensure that the cell cycle only progresses to its next phase if appropriate intra- and extracellular conditions are fulfilled. These conditions include the absence of DNA damage, proper alignment of the chromosomes in the metaphase plane, and abundance of nutrients and growth factors [1]. The sequential nature of cell cycle progression where the start of one phase depends on the completion of a previous phase resulted in the view of the cell cycle as a ‘domino-like’ process [2, 3].

Over the years, both theoretical and experimental studies have demonstrated how the mechanisms that control the ‘domino-like’ nature of cell cycle progression are centered around bistable switches. Such bistability is often encountered in systems that possess an S-shaped steady-state response curve and means that the system may settle in two different stable states depending on the initial conditions (Fig. 1A). These observable stable steady states correspond to the lower and top branch of the S-curve, while the middle part of the S-curve is unstable and cannot be measured experimentally. The full S-shaped response curve can therefore not be obtained experimentally, and bistability typically manifests itself through sharp jumps and hysteresis in the measurements. Hysteresis appears when the threshold for switching from low to high response levels is different from the threshold for switching from high to low response levels. In the cell cycle, these all-or-none responses ensure robust transitions between different cell cycle phases, while hysteresis prevents the cell cycle from returning to previous phases without having been through the whole cell cycle. Control mechanisms at checkpoints can prevent such transitions, either by keeping the input at sub-threshold levels or by shifting the right threshold to higher input levels [4–6].

**Fig 1.**
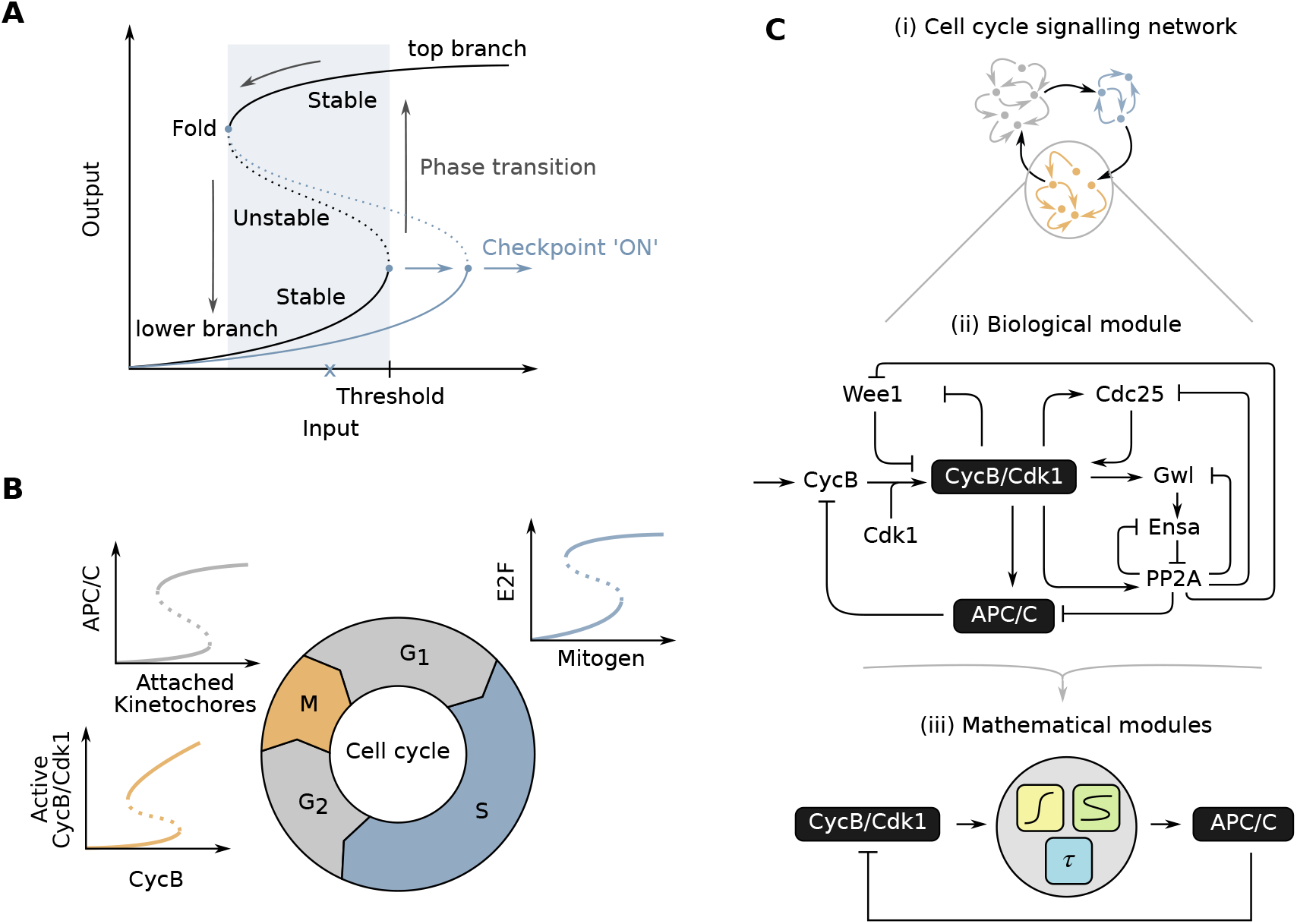
Modularity in the cell cycle. **(A)** An S-shaped response is characterized by the coexistence of two stable and one unstable steady states for a certain range of input levels (blue shaded area). Cell cycle transitions can be described as sudden jumps from the lower to upper branch of the curve (or vice versa). Checkpoints act by fixing the input at sub-threshold levels (x) or shifting the threshold value to a different input level (blue curve). **(B)** Different phase transitions in the cell cycle have been shown to be centered around bistable switches for which S-shaped response curves have been experimentally measured. **(C)** Entry into mitosis (and other cell cycle transitions) can be isolated as a biological module from the bigger network of other cell signaling events in the cell (i). Whereas the dynamical features of these modules have been studied extensively by considering detailed reaction schemes (ii), we propose a phenomenological approach based on three easily combinable ‘functional modules’: time delay, ultrasensitive and S-shaped responses (iii).

One of the first cell cycle transitions for which the role of an underlying bistable switch was established, was mitotic entry in early embryonic cells of *Xenopus* frogs. This cell cycle transition is characterized by the switch-like phosphorylation of numerous proteins, referred to as mitotic substrates. Throughout interphase, cyclin B (CycB) molecules are gradually produced and bind to cyclin-dependent kinase 1 (Cdk1). At the onset of mitosis, the phosphorylation state of these CycB-Cdk1 complexes abruptly changes, resulting in their switch-like activation and subsequent phosphorylation of the mitotic substrates [7]. The sudden changes in phosphorylation state of CycB-Cdk1 are generated by positive and double negative feedback loops with the phosphatase Cdc25 and kinase Wee1, respectively [8]. Theoretical work proposed that bistability, resulting from these feedback loops, plays an important role in cell cycle progression [9, 10]. Afterwards, this was experimentally validated [11–13]. Not only CycB-Cdk1, but also counteracting phosphatases such as PP2A help in regulating the phosphorylation state of mitotic substrates, and thus entry into and exit from mitosis. Recent findings showed how PP2A too is regulated in a bistable manner [14–16]. Furthermore, the phosphatases PP1 and Fcp1 have been implicated in regulating the exit from mitosis, but how exactly these phosphatases interact with the different cell cycle regulators remains a topic of active research [17, 18].

The observation that, just like M-phase entry, other phase transitions of the cell cycle are irreversible under normal physiological conditions, suggested that these too are built on bistable switches. For example, the transition from G1 to S phase is governed by a complex interplay between extracellular signals, CycD-Cdk4/6, the transcription factor E2F, retinoblastoma protein (Rb) and CycE-Cdk2. These interactions lead to bistability in the activity of E2F [19]. Appropriate conditions, such as a sufficient concentration of extracellular growth factors and nutrients, will push the cell across the threshold of the switch to a ‘high E2F’ state. At this point the cell irreversibly commits to the cell cycle, i.e. it will finish the started round of cell division even if nutrient levels drop [5, 20]. Another cell cycle transition for which bistability has been proposed to fulfill an important role is the metaphase-to-anaphase transition and the accompanying spindle assembly checkpoint (SAC). During this transition, microtubules in the mitotic spindle need to correctly attach to the sister chromatids. Once these are correctly attached, the cohesin rings that are keeping the sister chromatids together can be cleaved, upon which the chromatids are separated by the mitotic spindle. Although some experimental studies question the all-or-none nature of the SAC [21, 22], indirect experimental and theoretical findings support the idea that this transition is also centered around a bistable switch [23–26]. Given the recurring occurrence of all-or-none transitions throughout the cell cycle, the latter has been envisioned as a chain of interlinked bistable switches (Fig. 1B) [26, 27].

Although a chain of bistable switches can account for control mechanisms at cell cycle transitions and checkpoints, an additional mechanism is still needed to drive the cycle forward and reset it back to its initial state at the end — thus putting the dominoes back up after toppling them. This is provided by the periodic production and degradation of cyclins [28]. The CycB-Cdk1 complexes that accumulate during interphase are activated at mitotic entry. In turn, they will activate the Anaphase-Promoting Complex/Cyclosome (APC/C), a ubiquitin ligase. This protein complex then induces the degradation of the cyclins [29]. APC/C regulation is believed to be a complex multi-step mechanism in which time delays play an important role, hence introducing a delayed negative feedback loop in the reaction network [30, 31]. The latter is generally known to allow for robust and sustained oscillatory behavior of dynamical systems [32]. Even when the cellular checkpoints are absent, this negative feedback loop can drive autonomous biochemical oscillations in a ‘clock-like’ manner, as for example seen in *Xenopus* and sea urchin eggs which rapidly alternate between phases of DNA replication (S-phase) and segregation of the chromosomes (M-phase) [3, 33].

The cell cycle is a complex process with many interacting parts. Dividing it into a sequence of discrete modules such as bistable switches might seem artificial. However, since its introduction in biology by Hartwell *et al.,* [34], such a modular approach describing biological processes has been justified by advances in synthetic biology, genomics, cell signalling and single-cell techniques [35–38]. Furthermore, studying the cell cycle based on discrete modules is warranted by the temporal segregation of the different cell cycle phases, and the presence of bistability itself [39, 40].

Even though separating the full system into different modules greatly reduces the complexity, understanding the dynamical behavior of those modules often requires mathematical models instead of mere intuition [41]. Even for a single module, a biochemically detailed study results in a large number of variables and parameters, many of which are difficult to determine experimentally. As this might hamper interpretation of the results, it is often desirable to reduce the complexity of the mathematical model. One way to accomplish this is by focusing on a core subnetwork and omitting all other reactions, thus (hopefully) capturing the key qualitative behavior of the system [42]. Although this approach certainly has its merits, some dynamical properties such as bistability may be lost when reducing the model too much [43, 44]. Of note, this does not mean that models of small reaction networks imply less interesting dynamical features: for example, even a system containing a single molecule can behave in a bistable manner [45].

Another strategy to reduce the complexity of a model, while retaining much of the system’s dynamical behavior, is by making certain simplifying assumptions about the underlying reactions, rather than omitting them. For example, using timescale separation methods, one can identify variables that evolve on a much faster timescale than others. The equations for these variables can be replaced by expressions for the steady-state response curves, which can be introduced into the equations for the slow variables. This process is often responsible for the introduction of nonlinearities in a reaction system. Examples include Michaelis-Menten enzyme kinetics, zero-order ultrasensitivity, multisite phosphorylations, cooperative binding events and stoichiometric inhibition [46], whose net result is often summarized by their steady-state response curve. These steady state response curves can be described by reaction-specific formulas, such as the Michaelis-Menten equation for enzyme-kinetics [47] or Goldbeter-Koshland function for zero-order ultrasensitivity [48]. Alternatively, steep responses —of different molecular origins— can often be adequately approximated by a Hill equation [49–51]. The Hill equation is therefore used to introduce steep responses, without regarding the underlying reactions that generated them. Another example of dynamical behavior that can be explicitly incorporated are time delays, which arise in biochemical reactions due to the non-zero time to complete physical and/or biochemical processes, such as molecular transport or intermediate reactions [52, 53]. Such delays can be accounted for via delay differential equations. In addition to being simpler than mechanistic models, a phenomenological model based on explicit mathematical expressions is often the sole option to describe experimentally observed responses whenever the underlying molecular mechanisms are unknown.

S-shaped responses typically emerge from regulatory mechanisms like positive or double negative feedback loops [54]. Many cell cycles models include these feedback loops to generate the S-shaped response [10, 55–57]. Steep responses and time delays have been explicitly included in cell cycle models using a simple mathematical form. Given the recurring occurrence of bistable switches in the cell cycle, it is remarkable that such a direct mathematical formulation of S-shaped response curves has never been, as far as we know, explicitly incorporated into cell cycle models. Here we want to close this gap (Fig. 1C) by providing an easily tunable, phenomenological expression for such an S-shaped response. This expression can easily be converted into a functional module for ultrasensitivity by tuning a few parameters, and can be combined with a time delay. As such, we provide a toolbox of three functional modules (ultrasensitive response, S-shaped response, time delay) that can be combined in different configurations to model a variety of biological processes. We will illustrate this approach with models that include different bistable switches in the cell cycle.

## Results

### Ultrasensitive responses, S-shaped responses and delay as functional modules

Before considering actual models of the cell cycle, we start with a general overview of the functional modules that will be used in the rest of the paper: ultrasensitivity, S-shaped response and delay. Nonlinear response curves are often described by a Hill equation of the form 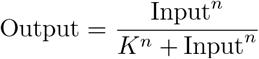. For *n* > 1, this gives a sigmoidal response curve, with higher values of *n* corresponding to steeper responses (Fig. 2A). Such steep responses are often said to be ‘ultrasensitive’ [51]. Our ultrasensitive module thus consists of a Hill function.

**Fig 2.**
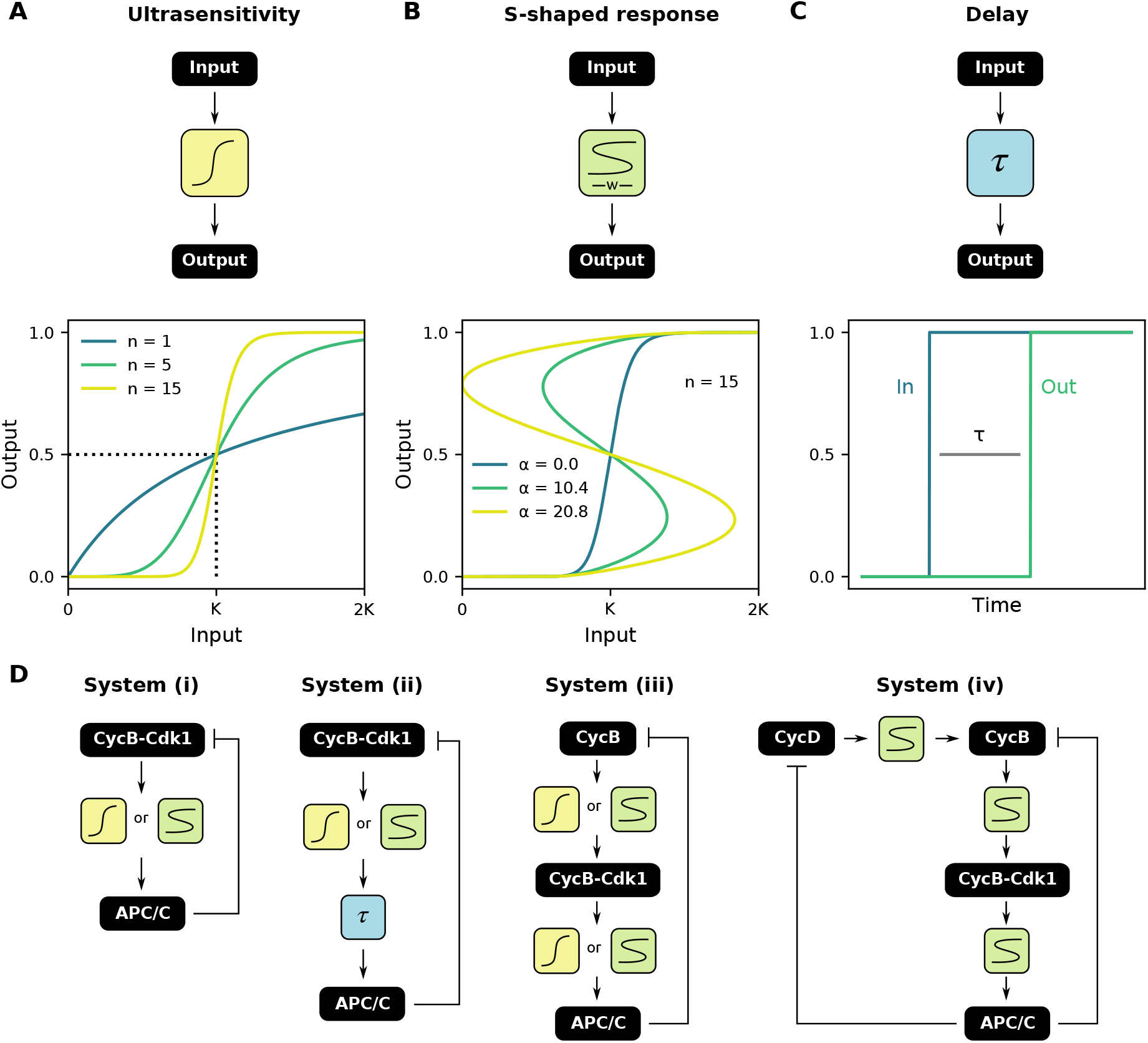
Ultrasensitive responses, S-shaped responses and delay as functional modules. **(A)** An ultrasensitive input-output response is characterized by the threshold value *K* and the Hill coefficient *n*. **(B)** An ultrasensitive function can be converted into a S-shaped response of different widths: for *α* = 0, an ultrasensitive response is obtained, while increasing *α* increases the width of the S-shaped region. **(C)** A certain lag time *τ* between input and output is encountered in several biochemical reactions. **(D)** Several combinations of the three functional modules, always in the presence of negative feedback, to describe oscillations in the cell cycle.

In order to produce a mathematical expression for our second module, the S-shaped response, we would like to ‘bend’ the ultrasensitive curve just described such that it becomes S-shaped (Fig. 2B). Mathematically, this can be achieved by starting from a Hill function, and multiplying the threshold value *K* with a scaling function *ξ*(Out):

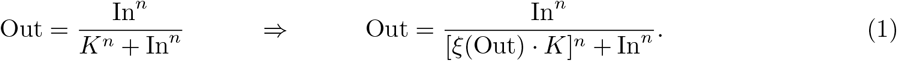

Note that the latter expression does not define a function — which is impossible, as per definition a S-shaped response has multiple outputs for one input. Instead, it should be interpreted as the steady state response of an ordinary differential equation (ODE):

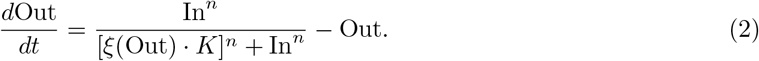

Although different options for *ξ* are possible, a straightforward choice in analogy with, for example, the Fitzhugh-Nagumo model would be a cubic function [58]:

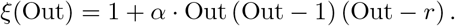

Changing the parameter *α* allows for a smooth transition between an ultrasensitive and S-shaped response: for *α* = 0 we retrieve the original ultrasensitive response, while increasing its value results in wider S-shaped regions (Fig. 2B). The parameter *r* can be used to make the response curve asymmetric. In the remainder of the text, we keep *r* = 0.5 (as in Fig. 2B), and we will not interpret the parameter *r* further biologically. The effect of the different parameters on the shape of the scaling function is discussed in more detail in S1 Text and S1 Fig.

For the last module, i.e. delay, a lag time *τ* between input and output of the system (Fig. 2C) can be explicitly modeled by setting Out(*t*) = In(*t – τ*). This turns the equation into a delay differential equation. In what follows, we will combine these three functional modules in several configurations, always in combination with a negative feedback (Fig. 2D), to model different aspects of the cell cycle. We will explain how changing the mathematical properties and function parameters influence the overall dynamical behavior of the model.

### S-shaped, but not ultrasensitive, responses cause a two-dimensional system of the cell cycle to oscillate

The most straightforward strategy for combining the different functional modules into actual cell cycle models, would be to start with a simple — but biologically relevant — reaction network and gradually add additional modules to increase the scope of the model. The simplest cell cycle models arguably describe the ‘clock-like’ cell cycle oscillations in *Xenopus laevis* eggs, compared with the more complex ‘domino-like’ mechanism in somatic cells. Indeed, the early embryonic cell cycle in *X. laevis* simply cycles between S- and M-phase and lacks checkpoints and gap phases [33]. Furthermore, cell cycles 2 till 12 after fertilization of the egg are characterized by an increased activity of the phosphatase Cdc25 relative to the kinase Wee1, resulting in quick activation of CycB-Cdk1 and subsequent APC/C activation [59]. Taken together, the reaction network at the core of the early embryonic cell cycle can be simplified by a negative feedback loop, where active CycB-Cdk1 phosphorylates and activates APC/C, which then causes degradation of CycB molecules. A two-variable phenomenological model of this system is given by (Fig. 3A):

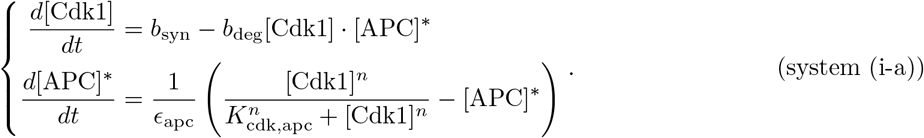

**Fig 3.**
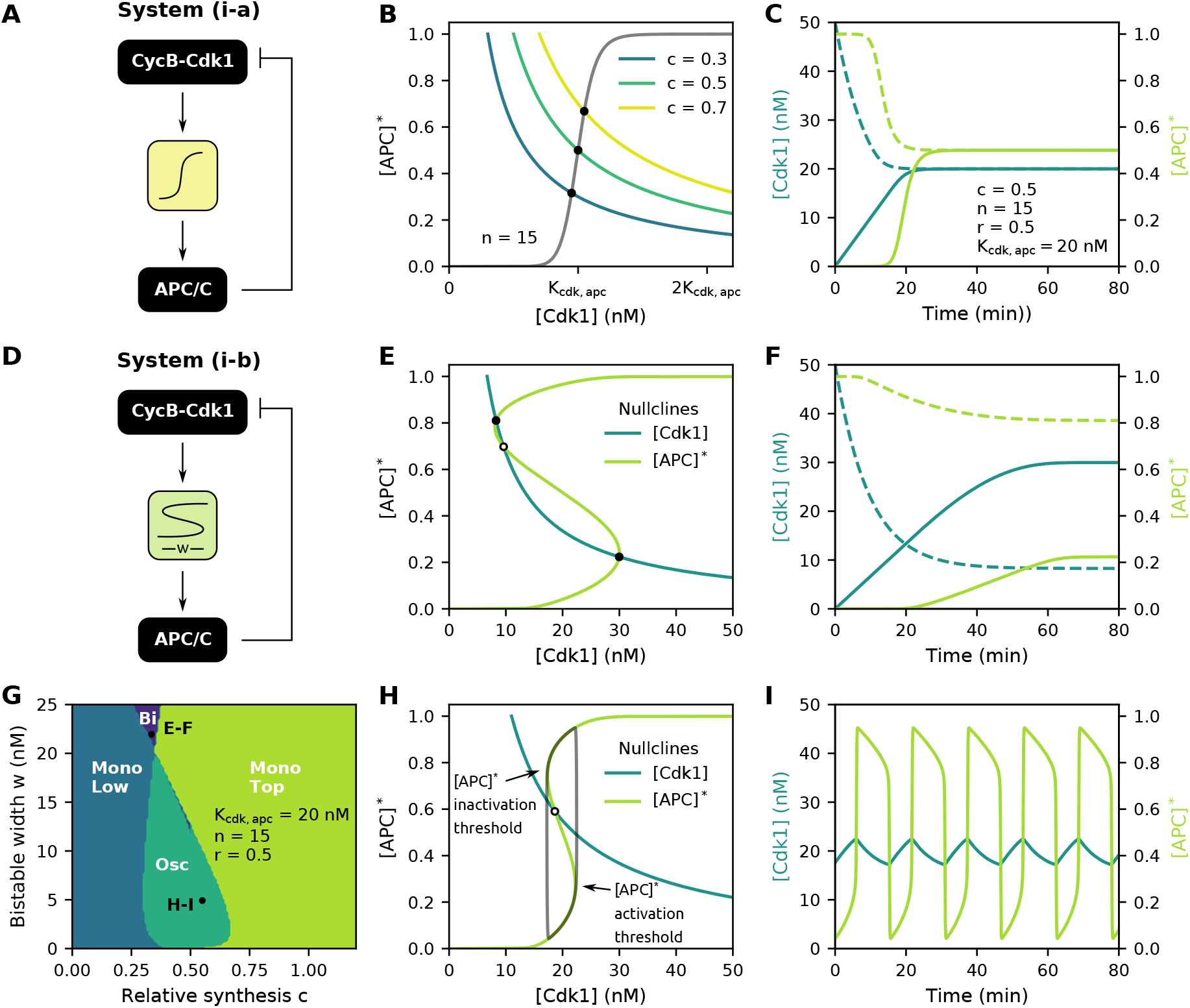
S-shaped, but not ultrasensitive, responses cause a two-dimensional system of the cell cycle to oscillate. **(A)** Block diagram of the ultrasensitive, negative feedback system (i-a). **(B)** Effect of relative synthesis *c* on the location of the [Cdk1] nullcline in the phase plane. **(C)** Time traces for two different initial conditions of system (i-a), denoted by either continuous or dotted line. Only one steady state exists. **(D)** Block diagram of the bistable, negative feedback system (i-b). **(E)** For wide bistable [APC]* nullclines, three intersections in the phase plane can exist, resulting in bistability of the overall ODE system. Closed black dots depict stable steady states, while open dots denote unstable steady states. **(F)** Time traces for system (i-b). Depending on the initial conditions, the system will settle in one of two stable steady states. **(G)** Number and position of the steady states for system (i-b). Osc = oscillations, Bi = bistable, Mono Low = one stable steady-state on lower branch of [APC]* nullcline, Mono Top = one stable steady-state on top branch of [APC]* nullcline. **(H-I)** If the [Cdk1] nullcline intersects the [APC]* nullcline in between its folds, one unstable steady state exists, resulting in oscillations. The used parameter values can be found in the panels.

Here, the two variables [Cdk1] and [APC]* represent the concentrations of activated CycB-Cdk1 complexes and the ratio of activated APC/C to total APC/C, respectively. The first equation describes the rate of change of activated CycB-Cdk1 complexes and consists of two terms: the constant synthesis of cyclins and their APC/C-dependent degradation. In this model all synthesized cyclin immediately binds to Cdk1 and activates it. This is why cyclin synthesis is directly included in the equation for [Cdk1]. The second equation in system (i-a) states that the rate of change of [APC]* is proportional to the difference between the experimentally determined [APC]* ultrasensitive steady state response and its current level (similar as in Eq. 2) [30, 59], which it approaches at a time scale 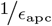.

Because system (i-a) has two variables, its behavior can be conveniently studied in the phase plane. To determine the steady state behavior of system (i-a), one needs to find the number and location of intersections of the [Cdk1] and [APC]* nullclines. These are the curves in the [Cdk1]-[APC]* phase plane for which 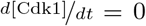 and 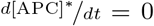 respectively. Here, only one intersection can exist, whose location is determined by the relative position of both nullclines (Fig. 3B-C). Moreover, this steady state is linearly stable (see S1 Text for details). While parameters *n* and *K*_cdk,apc_ affect the ultrasensitive [APC]* nullcline, the location of the [Cdk1] nullcline is determined by the ratio of *b*_syn_ and *b*_deg_. However, after non-dimensionalizing the system to facilitate mathematical analysis (see Methods for details), this ratio can be re-expressed as the product of *K*_cdk,apc_ with a newly introduced dimensionless parameter *c*, termed the relative synthesis rate:

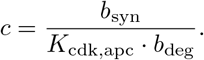

Changing *c* thus changes the location of the [Cdk1] nullcline in the phase plane: whereas low values shift it to the left, high *c* values shift the [Cdk1] nullcline to the right (Fig. 3B). It is mainly this parameter that will be used in the remainder of the text when assessing the effect of the [Cdk1] nullcline on the dynamical behavior of the overall system.

As the steady state is stable, this system can not serve as a basic model to describe cell cycle oscillations. To make the system oscillate, either a time delay or a S-shaped response can be introduced. The effect of a time delay in combination with ultrasensitivity has previously been studied in detail [60] (see also S2 Fig). Here we will focus on the effect of converting the ultrasensitive module from system (i-a) into a S-shaped module, which is in line with recent experimental findings that measured hysteresis in APC/C response curves [15]. As discussed above, the threshold *K*_cdk,apc_ can be multiplied with a cubic scaling function *ξ*([APC]^*^) = 1 + *α*_apc_[APC]*([APC]* − 1)([APC]* − *r*) (Fig. 3D):

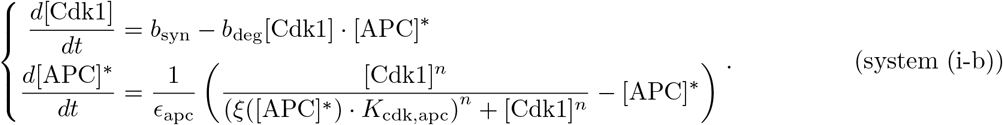

Note that by multiplying the threshold *K*_cdk,apc_ with *ξ* in system (i-b), the [APC]* nullcline becomes S-shaped when *α*_apc_ *>* 0 (as introduced in Eq. 2 and Fig. 2B). Systems exhibiting such a S-shaped response are good candidates for showing bistability, in which case the system can evolve over time to two steady states. Whether this behavior is indeed observed, depends on the number of intersections between the [Cdk1] and [APC]* nullclines in the two-dimensional phase plane. As the [APC]* nullcline is now S-shaped instead of sigmoidal, either one or three intersections can exist with the [Cdk1] nullcline (Fig. 3E,G,H). The system is bistable when three intersections exist, with one steady state being unstable and the other two stable. It is the initial condition of the system that determines in which of the two stable ones the system will settle (Fig. 3F). Whenever the system has only one steady state (i.e. one intersection of the nullclines), the latter can either be stable (on the upper or lower branch of the S-shaped nullcline) or unstable. Only in the latter case, i.e. if both nullclines intersect in between the fold points of the [APC]* nullcline (Fig. 3H), sustained oscillations around this steady state appear. These oscillations can be thought of as ‘orbiting around the nullclines’, i.e. they show increasing [Cdk1] levels on the lower branch of the response curve until the [APC]* activation threshold is reached. At this point, [APC]* levels jump to a higher value, which triggers the decrease of [Cdk1]. The system then proceeds along the upper branch until the inactivation threshold is reached and the cycle is complete. These oscillations, characterized by slow progress along the branches of the nullclines and quick jumps between them, are called relaxation oscillations. They are similar to oscillations observed in, for example, the FitzHugh-Nagumo equations [58].

From the observation that the system oscillates if the nullclines have a single intersection in the middle, we can derive some qualitative conditions for oscillations to exist: the [APC]* nullcline should not be too wide and the [Cdk1] nullcline should not be located too far to the left nor to the right (relative to the other nullcline). Indeed, from Fig 3G, we see that oscillations are favored for *c* values around 0.5 and narrow S-shaped regions. As the S-shaped region becomes wider, the period of oscillations and [Cdk1] amplitude increases (S3 Fig), but the range of relative synthesis *c* for which oscillations can be sustained decreases. For [APC]*, the effect of the width of the S-shaped region on the amplitude is more moderate, and the amplitude is near-maximal for most oscillations. From the time profiles, we further see that the overall shape of the oscillations has a typical sawtooth-like waveform (Fig. 3I). It is interesting to note that, in this system, the effect of the S-shaped response is mixed: on the one hand, it is required for the system to oscillate, but on the other hand a wider S-shaped region makes oscillations less likely, as in that case the system settles in a steady state on the upper or lower branch.

### Asymmetric S-shaped APC/C nullclines slow down APC/C (in)activation

The use of a cubic function *ξ* with *r* = 0.5 results in what we would call ‘symmetric’ S-shaped responses, meaning that the vertical distance between the left folding point and lower branch equals the vertical distance between the right folding point and the upper branch of the response curve (S4 Fig). One might then wonder what would happen if the assumption of such symmetry is relaxed. After all, the lower and top branch of experimentally observed S-shaped response curves (recall that the middle part cannot be measured experimentally) will typically not be symmetric. Examining some of the experimentally measured responses in [15], such as the one for APC/C phosphorylation levels, indeed shows that asymmetry can occur.

To explore the effect of asymmetry, one needs to shift the folding points of the S-shaped response up or down. Using the cubic *ξ* function, this can be accomplished by altering the value of parameter *r*. However, this approach would shift the response curve horizontally too, making it impossible to isolate the effect of vertical asymmetry from other effects. Similar difficulties exist when *ξ* is defined by other smooth functions, such as a quadratic function (S4 Fig). As a solution to this problem, we approximated the cubic scaling function by a piecewise linear function in combination with large values of *n* (*n* = 300). The piecewise approach enables us to exactly position the extrema of *ξ*([APC]*) at the desired location, while the high values of *n* ensure that these extrema correspond to the folds of the S-shaped nullcline (see S1 Text for further details), thus giving precise control over their location.

The approximation of the cubic scaling function *ξ*([APC]*) (with arbitrarily chosen values for *r* and *α*_apc_) by a piecewise fit of linear functions through its extrema (inset Fig. 4A), leads to the response curve in Fig. 4A. When comparing the dynamics of the system with the piecewise approximation to the dynamics with a smooth cubic function, similar oscillatory regions in parameter space can be observed (Fig. 4B). We then exploited the fact that we can now independently change the [APC]* coordinate of the local extrema of *ξ*([APC]*) to explore the effect of an asymmetric response curve: by moving the extrema of *ξ*([APC]*) to the right or left, the corresponding folds of the S-shaped response shifted up or down in the phase plane respectively, while their [Cdk1] coordinate was kept constant at the levels from Fig 4A. Again, the system can be either bistable or monostable, or have an unstable steady state leading to oscillations. The period of these oscillations decreases as the vertical distance between the folds increases, i.e. towards the lower right corner of Fig 4C.

**Fig 4.**
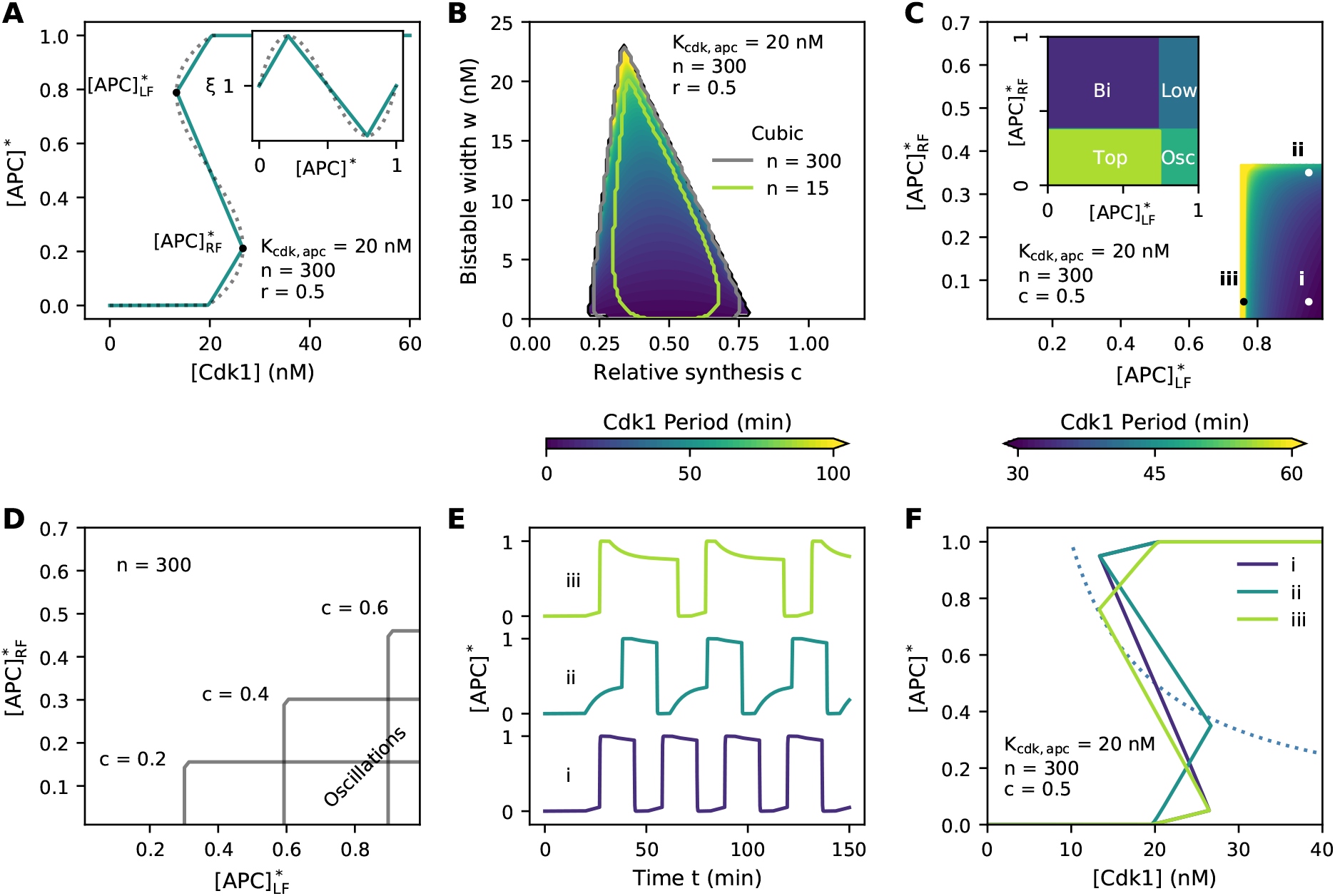
Asymmetric bistable APC/C nullclines slow down APC/C (in)activation. **(A)** Piecewise approximation of cubic scaling function *ξ*([APC]*) (inset) and derived bistable response curve. **(B)** Comparison of oscillatory region for large *n* between a cubic *ξ*([APC]*) (grey contourline) and its piecewise approximation (colormap). The green contour line indicates the oscillatory region for a cubic scaling function with *n* = 15. **(C)** [Cdk1] period for different [APC]* coordinates of the folds ([Cdk1] coordinates were kept constant at the levels from panel A). Inset: number and location of steady-states. **(D)** Oscillatory regions for different values of c. **(E,F)** Time traces and phase plane for parameter values indicated in panel C. The used parameter values can be found in the panels.

The position of the [Cdk1] nullcline (set by the parameter *c*) determines the positions of the fold points for which the system can oscillate. For smaller *c* values, oscillations can occur for low left folds, while these need to be higher as the [Cdk1] nullcline shifts to the right (higher *c*). The further apart the folds, the more likely the [Cdk1] nullcline intersects the [APC]* nullcline in between the folds. This can be seen from the overlap of the oscillatory region for all *c* values in the lower right corner of the parameter space (Fig. 4D).

The vertical position of the folds also determines the shape of the time traces of [APC]* activity. For symmetric nullclines, here meaning [APC]*_*RF*_ = 1 − [APC]*_*LF*_, with vertically well-separated folds, the time trace resembles a square wave (Fig. 4E,F i). This leads to rapid and abrupt interconversion between active and inactive [APC]*, leading to well-defined transitions between interphase and M-phase. Shifting the right fold upwards generates an asymmetric nullcline and introduces a period of slow exponential-like build-up in [APC]* activity before it switches to its maximal activity (Fig. 4E,F ii). Hence, the activation of [APC]* is slowed down as the time needed to convert completely inactive [APC]* into its fully activated form is lengthened. Likewise, shifting the left fold downwards results in a slow exponential-like decay in [APC]* activity prior to complete inactivation and thus in a longer time required to completely inactivate all [APC]* (Fig. 4E,F iii).

### The S-shaped module reproduces the behavior of mass-action models

In the previous sections, we showed how the S-shaped module can be used to conveniently model oscillatory systems and showed that the obtained oscillations resemble those of the early embryonic cell cycle in a qualitative way. The way we introduced the S-shaped response, through the modified Hill function, is artificial, and not based on biochemical interactions. To show that our approach can represent the behavior of an actual biochemical system, we compare our module-based system to an existing cell cycle model based on mass-action kinetics. For this purpose, we first extended a previously described mass-action model of the PP2A-ENSA-GWL network [61], by incorporating synthesis and APC/C-mediated degradation of CycB (Fig. 5A). This system is known to generate S-shaped APC/C response curves via a double negative feedback loop. More specifically, GWL (which is phosphorylated by CycB-Cdk1) indirectly inhibits PP2A by phosphorylating ENSA, which is both substrate and inhibitor of PP2A [62]. PP2A itself, when active, dephosphorylates and inactivates GWL, thus closing the double negative feedback loop. This leads to an S-shaped response of PP2A activity as function of CycB-Cdk1 activity. As APC/C is a substrate of PP2A, the S-shaped steady-state response of PP2A is translated into a similar response for APC/C as function of CycB-Cdk1.

**Fig 5.**
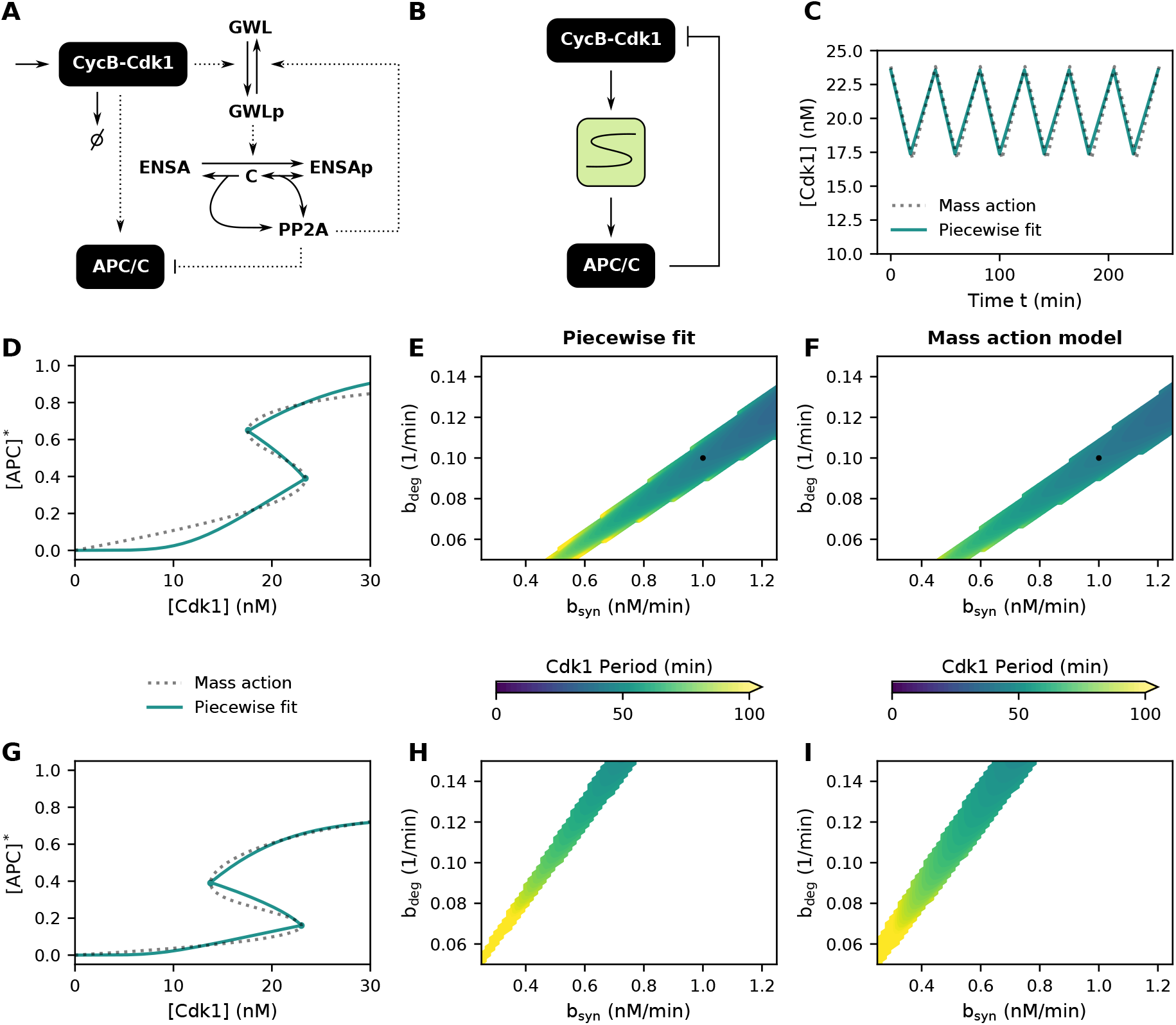
The S-shaped module reproduces the behavior of mass-action models. **(A)** Mass-action model of the PP2A-ENSA-GWL network in which double negative feedback can lead to S-shaped response of APC/C. **(B)** The S-shaped module replacing the detailed reaction network in panel (A). **(C)** Time traces showing the resemblance between the modular approach and mass-action model. Parameters as in panel (D), with synthesis and degradation as indicated in (E,F). **(D)** Steady state response curve for mass-action model and fitted S-shaped module. Parameter values as given in Table 1, association constant of ENSAp with PP2A *k*_ass_ = 5.62 nM^−1^min^−1^ and APC/C dephosphorylation constant *k*_da_ = 3.16 nM^−1^min^−1^. **(E,F)** Oscillatory region as a function of CycB synthesis and degradation rates for the piecewise fit and mass-action model. Black dot denotes rate constants used in panel (C). **(G,H,I)** Similar as panels (D,E,F) with larger association constant (*k*_ass_ = 7.94 nM^−1^min^−1^), making the S-shaped region wider, and larger APC/C dephosphorylation constant (*k*_da_ = 10 nM^−1^min^−1^), shifting the response curve downwards.

Once the S-shaped steady-state response curve of the mass-action model (for a fixed parameter set) was calculated, we approximated this response using a piecewise linear scaling function *ξ*([APC]*) (Fig. 5D, see Methods for details). Next, we screened different values for the synthesis and degradation rate of CycB for both modeling frameworks (note how these parameters do not affect the shape of the response curve itself, but only alter the way the time series orbit around it). Even though the steady-state response curves do not exactly match, the region in parameter space for which oscillations can be observed do correspond to a good extent. Furthermore, the periods of the oscillations for both models are very similar, as can also be seen from the time traces in Fig. 5C. This behavior is not unique to the chosen parameter set, as also other response curves and their corresponding oscillations can be approximated well (Fig. 5G-I). Hence, the S-shaped module can faithfully represent the dynamical behavior of more detailed mass-action models.

**Table 1.** Default parameter values for the mass-action model of the PP2A-ENSA-GWL network.

| Parameter | Value | Units | Description |
| --- | --- | --- | --- |
| $k_{\text{pg}}$ | 0.03 | 1/(nM min) | Phosphorylation constant of GWL |
| $k_{\text{dg}}$ | 3.16 | 1/(nM min) | Dephosphorylation constant of GWL |
| $k_{\text{pe}}$ | 3.16 | 1/(nM min) | Phosphorylation constant of ENSA |
| $k_{\text{pa}}$ | 0.63 | 1/(nM min) | Phosphorylation constant of APC/C |
| $k_{\text{dis}}$ | 1 | 1/min | Dissociation constant of ENSA-PP2A complex C |
| $k_{\text{cat}}$ | 3.98 | 1/min | Catalytic constant of ENSA-PP2A complex C |
| $\text{GWL}_{\text{tot}}$ | 20 | nM | Total GWL concentration |
| $\text{ENSA}_{\text{tot}}$ | 40 | nM | Total ENSA concentration |
| $\text{PP2A}_{\text{tot}}$ | 20 | nM | Total PP2A concentration |

### Delay increases the period, amplitude and robustness of oscillations

So far, we have described the regulation of APC/C by CycB-Cdk1 as a module in which changes in CycB-Cdk1 immediately affect APC/C activity. In reality, however, this regulation is believed to be a complex multi-step mechanism in which time delays play an important role [30, 60]. Two types of delays can be distinguished. First, a delay *τ*_1_ between the activation of CycB-Cdk1 and the subsequent activation of APC/C can exist, i.e. a lag time between [Cdk1] levels reaching the right fold of the [APC]* nullcline and [APC]* levels actually jumping to the upper branch (Fig. 6C-F). Secondly, a delay *τ*_2_ might occur between the inactivation of CycB-Cdk1 and APC/C, respectively. The delays *τ*_1_ and *τ*_2_ do not necessarily need to be the same. For example, if the activation of APC/C were to take much longer than its inactivation, delay *τ*_1_ would have a larger value than *τ*_2_ [60]. This time delay can be included directly into the model equations (Fig. 6A):

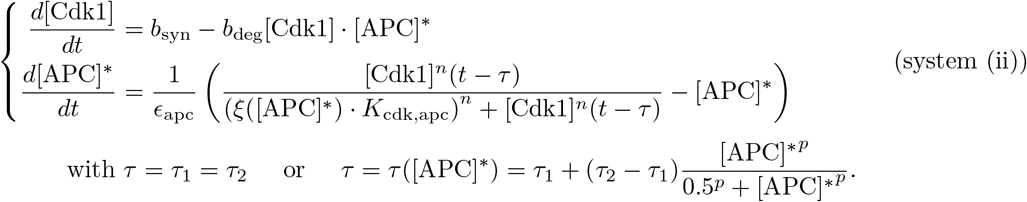

**Fig 6.**
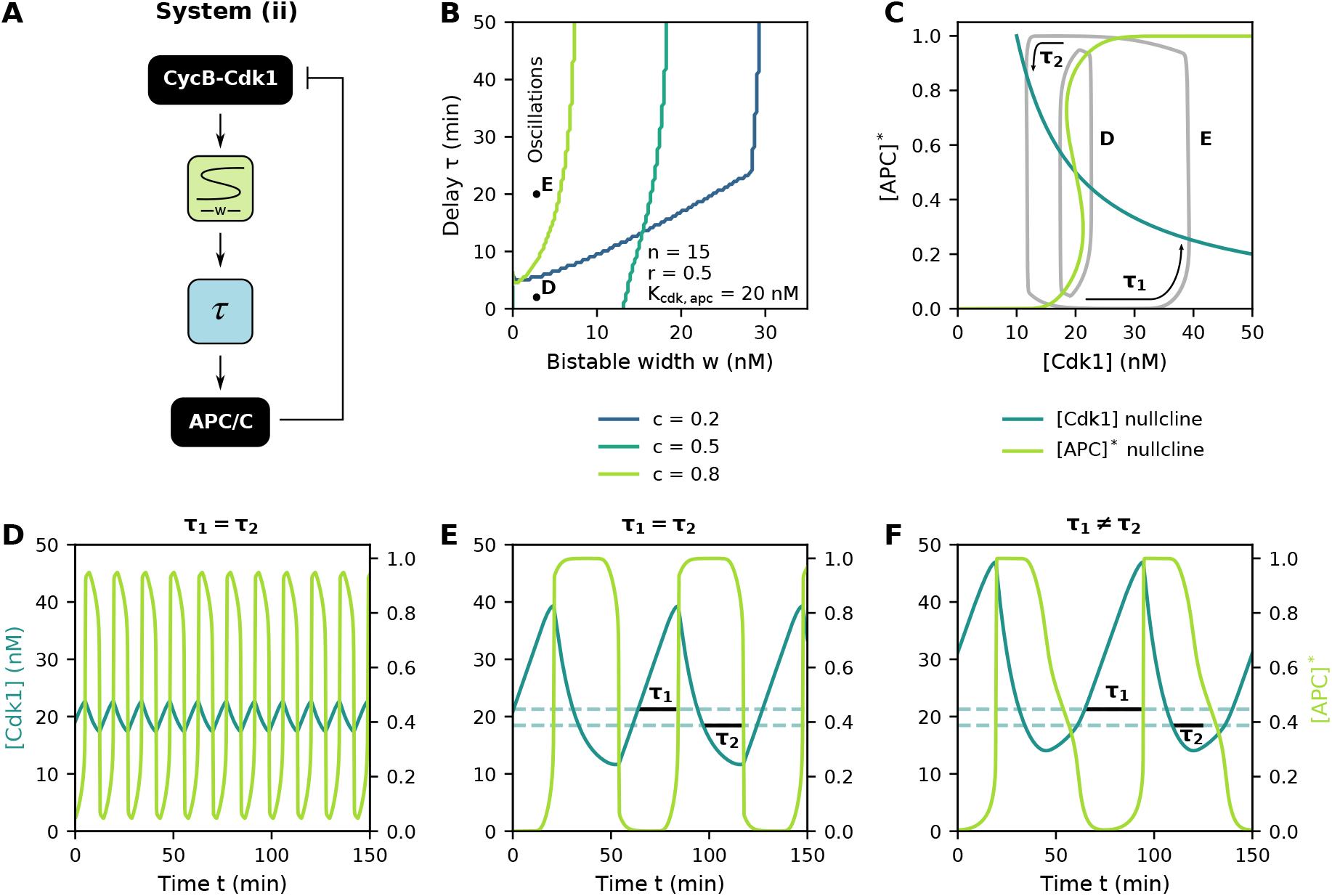
Delay increases the period, amplitude and robustness of oscillations. **(A)** Block diagram of the bistable, delayed negative feedback network. **(B)** Oscillatory regions, located above and to the left of the plotted lines, as a function of the time delay and bistable width for different values of the relative synthesis rate *c*. See figure for parameter values. **(C)** Phase plane representation of the oscillations for different values of *τ*. Note how one can only conclude that *τ*_1_ = *τ*_2_ from time traces, not directly from the phase plane. **(D,E)** Time traces for parameter values indicated in panel B. *τ*_1_ denotes the delay between activation of [Cdk1] and activation of [APC]*, while *τ*_2_ represents the delay between their inactivations. Dashed lines depict the [Cdk1] levels at the folds of the bistable [APC]* nullcline. **(F)** State dependent time delays can account for differences in the activation and inactivation delays (*p* = 5).

The expression for *τ*([APC]*) can be used in case *τ*_1_ ≠ *τ*_2_, as then the value of the delay is not a constant but depends on the level of [APC]* in the system. Here, we use a Hill equation in *τ*([APC]*) to ensure that the time delay is a smooth function of [APC]*. However, for high values of the Hill coefficient *p*, the Hill function approaches a step function with *τ ≈ τ*_1_ if [APC]* is smaller than 0.5 and *τ ≈ τ*_2_ when [APC]* activity is larger than 0.5, the situation we want to capture with the model.

For *ξ*([APC]*) = 1 (i.e. *α*_apc_ = 0) we retrieve the combination of a delay and ultrasensitive module (S2 Fig). As before, increasing *α*_apc_ converts the ultrasensitive response into a S-shaped one. Without a time delay, i.e. *τ* = 0, system (ii) is identical to system (i-b): oscillations can be observed for intermediate values of the relative synthesis *c*(*c* = 0.5) whenever the S-shaped region is not too wide (as otherwise the overall system becomes bistable). Low or high *c* values result in one stable steady state and no oscillations (compare Fig. 6B and Fig. 3G). However, adding a sufficiently large time delay *τ* can cause the system to start oscillating, even for widths of the S-shaped region and values of *c* where the system is mono- or bistable if *τ* = 0. The wider the S-shaped region becomes, the larger the delay required to induce oscillations. A maximal width exists beyond which no oscillations can be sustained, independent of the time delay, corresponding to the right vertical boundary of the oscillatory region in Fig 6B (especially clear for *c* = 0.2).

Both the period and [Cdk1] amplitude grow as the time delay increases (Fig. 6C-E, S5 Fig). Activation of [APC]*, on the other hand, is an all-or-none process for the majority of parameter values, with longer delays prolonging the time during which the [APC]* activity is at its maximal/minimal level, resulting in plateau phases in the time profiles (Fig. 6C-E, S5 Fig).

### Large amplitude oscillations are facilitated by two bistable switches

In the previous section we considered CycB-Cdk1 activity to be equivalent to CycB levels. This is justified for cycles 2-12 of the embryonic cell cycle of *X. laevis*, where all CycB quickly binds to Cdk1 and the complexes are rapidly activated by the phosphatase Cdc25 whose activity is dominant over the inhibitory kinase Wee1 [59]. However, in the first embryonic cycle and extracts of *Xenopus* eggs, a S-shaped response of CycB-Cdk1 activity as function of total CycB concentrations is typically observed [11, 12, 59]. Hence, a logical next step in adjusting our cell cycle model is to add a separate S-shaped module to account for this regulation (Fig. 7A):

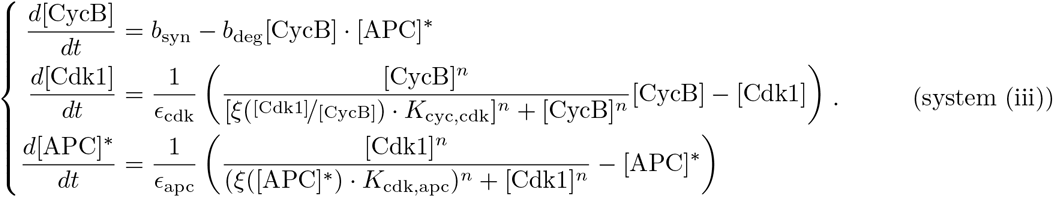

**Fig 7.**
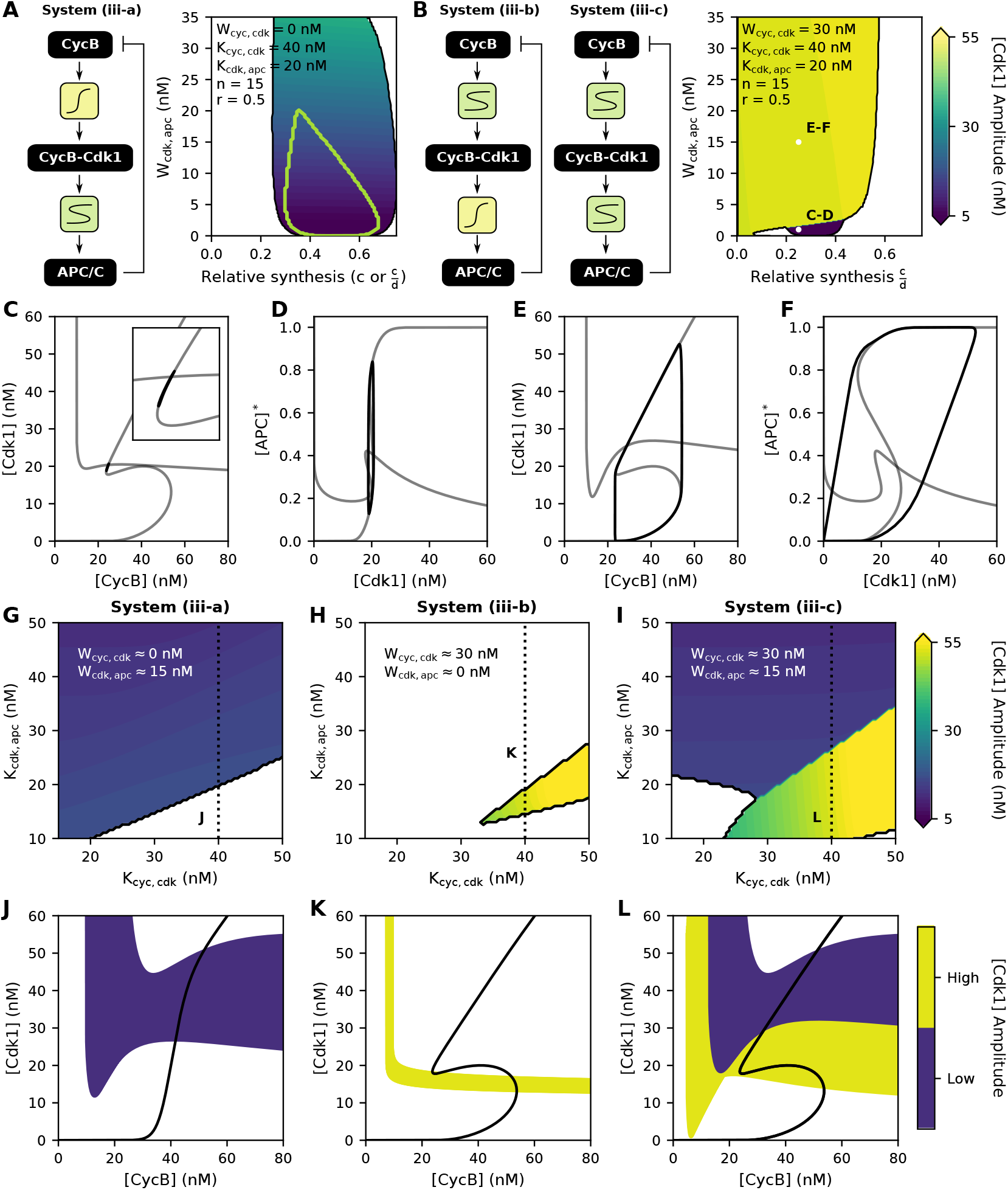
Large amplitude oscillations are facilitated by two bistable switches. See figure for parameter values. **(A)** [Cdk1] amplitude for system (iii-a), with relative synthesis 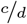. The green contour represents the oscillatory region for system (i), with relative synthesis *c*. **(B)** [Cdk1] amplitude for system (iii-b) and (iii-c). **(C-F)** Time trajectories (black lines) orbiting around the nullclines (grey lines) showing the distinct features of low and high amplitude oscillations for parameters as shown in panel B. **(G-I)** [Cdk1] amplitudes for the different system configurations as a function of the threshold values *K*. Parameters as in panel B or as indicated, with *c* = 0.5. **(J-L)** Phase plane representation of the oscillatory regions in G-I, as indicated by the dotted lines.

Here, the variable [CycB] represents the total concentration of CycB-Cdk1 complexes, [Cdk1] represents activated CycB-Cdk1 complexes and [APC]* the ratio of active APC/C molecules to the total amount of APC/C. We can still consider the total amount of CycB-Cdk1 (but not its activity) to be equivalent to the total amount of CycB, as Cdk1 is present in excess and it is assumed that free CycB molecules immediately bind free Cdk1 molecules [63]. Note how for the rate of change of [Cdk1], the Hill-like expression for the S-shaped response is multiplied by [CycB], the reason for which is explained in detail in the Methods.

Adding this second module for CycB-Cdk1 regulation greatly affects the position and shape of the nullclines, which now become surfaces in the three dimensional space (although they can be projected in a 2D plane, Fig. 7C-F). As a result, the parameter space sustaining oscillations changes compared with system (i-b), even if the added response of [Cdk1] as a function of [CycB] is only ultrasensitive, as in system (iii-a) which can be derived from system (iii) by setting the prefactor *α*_cdk_ in 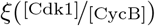 equal to zero (Fig. 7A). In particular, oscillations can be observed for wider [APC]* nullclines. Note that whereas the relative synthesis (i.e. 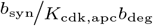) was given by the previously introduced parameter *c* for system (i-b), the non-dimensionalization of system (iii) (see Methods for details) causes the relative synthesis to become the ratio of *c* and a newly introduced parameter 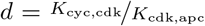. The [Cdk1] amplitude is directly proportional to the width of the [APC]* nullcline (Fig. 7A and S6 Fig), and the amplitude is minimally affected when moving the nullclines in the phase plane by altering their threshold values *K* (Fig. 7G,J and S6 Fig).

Whenever the [Cdk1] response as a function of [CycB] becomes bistable by making *α*_cdk_ in system (iii) greater than zero, the behavior of the system drastically changes, with two distinct types of oscillations emerging corresponding to either low or high [Cdk1] amplitudes (Fig. 7B, system (iii-b) for *W*_*cdk,apc*_ = 0 and (iii-c) for *W*_*cdk,apc*_ > 0). Low amplitude oscillations arise from trajectories orbiting around the [APC]* nullcline, but moving along the linear part of the [Cdk1] nullcline (Fig. 7C-D). Consequently, their amplitude is determined by the width of the [APC]* nullcline, similar as when the [Cdk1] nullcline was ultrasensitive. High amplitude oscillations, on the other hand, do orbit around the [Cdk1] nullcline (Fig. 7E-F), the linear part of which then becomes instructive for the amplitude of the oscillations. Therefore, the amplitude will increase if the [Cdk1] nullcline is shifted to higher threshold levels *K*_cyc,cdk_ (Fig. 7H-I and S6 Fig).

The high amplitude oscillations generated by two bistable switches induce a sufficiently large increase in CycB-Cdk1 activity followed by sufficiently large inactivation required for correct entry into and exit from mitosis. Furthermore, the two bistable switches make such oscillations more robust, meaning that oscillations can be observed for a larger set of parameter values. From a biological perspective, this means that cell cycle oscillations can be ensured even if physiological conditions within the cell (e.g. enzyme activity) would fluctuate, a situation which might otherwise disrupt correct cell cycle progression. This behavior can be appreciated from the larger yellow region in Fig. 7I,L compared with Fig. 7H,K. Note that the nullclines need to intersect in between the folds of the S-shaped response in system (iii-b) (Fig. 7K) and the previously considered two-dimensional system (i) to get oscillations, whereas this is not the case for system (iii-c) with two bistable switches (Fig. 7L). Here, oscillations can be observed even if the second nullcline intersects the [Cdk1] nullcline at its linear part, as long as this second nullcline reaches [Cdk1] values lower than the left fold of the [Cdk1] nullcline.

### The cell cycle can be represented as a chain of interlinked bistable switches

The results on the G2-M transition coupled to a (delayed) negative feedback in the previous sections are mainly applicable to the early embryonic cell cycles that rapidly cycle between S- and M-phase without intermittent gap phases. Many insects, amphibians and fish that lay their eggs externally carry out multiple rounds of such rapid cell cycles following fertilization. However, all of them then pass through the so-called midblastula transition (MBT) after approximately ten cell cycles [64–67]. This transition is characterized by the establishment of gap phases and slowing down of the cell cycle, resulting in a higher resemblance to the cell cycle as typically studied in yeast and mammalian somatic cells. Remarkably, many of the transitions between these additional cell cycle phases are — just like the G2-M transition — governed by bistable switches [26].

During the G1-S transition in cultured mammalian cells for example, it is the activity of the E2F transcription factor that is regulated in a bistable manner. The external presence of growth factors can induce the expression of CycD and the activation of CycD-Cdk4/6 complexes, followed by the activation of E2F. E2F then induces the expression of several target genes, one of which is CycE. As CycE can further enhance the activation of E2F, a positive feedback loop is established, ultimately leading to the bistable response of E2F [19]. Consequently, the G1-S transition can be depicted as a S-shaped module, similar as was done for the G2-M transition.

It is possible to combine the different transitions — each represented by a S-shaped module — into a phenomenological model of the overall cell cycle. For this, we still need to link the ‘input’ and ‘output’ of each module in a suitable way. We link E2F activity (the output of the G1-S switch) to CycB (the input of the G2-M switch) by recognizing that CycA, which is a target of E2F, drives the activation of the transcription factor FoxM1, which subsequently induces the expression of CycB [68] (Fig 8A, see Methods for associated equations). Similar as before, cyclins then need to be degraded to reset the system to the lower branch of the S-shaped response curves, after which a new round of cell division can start. Here, it is assumed that the degradation of all cyclins is induced in M-phase by activated APC/C.

**Fig 8.**
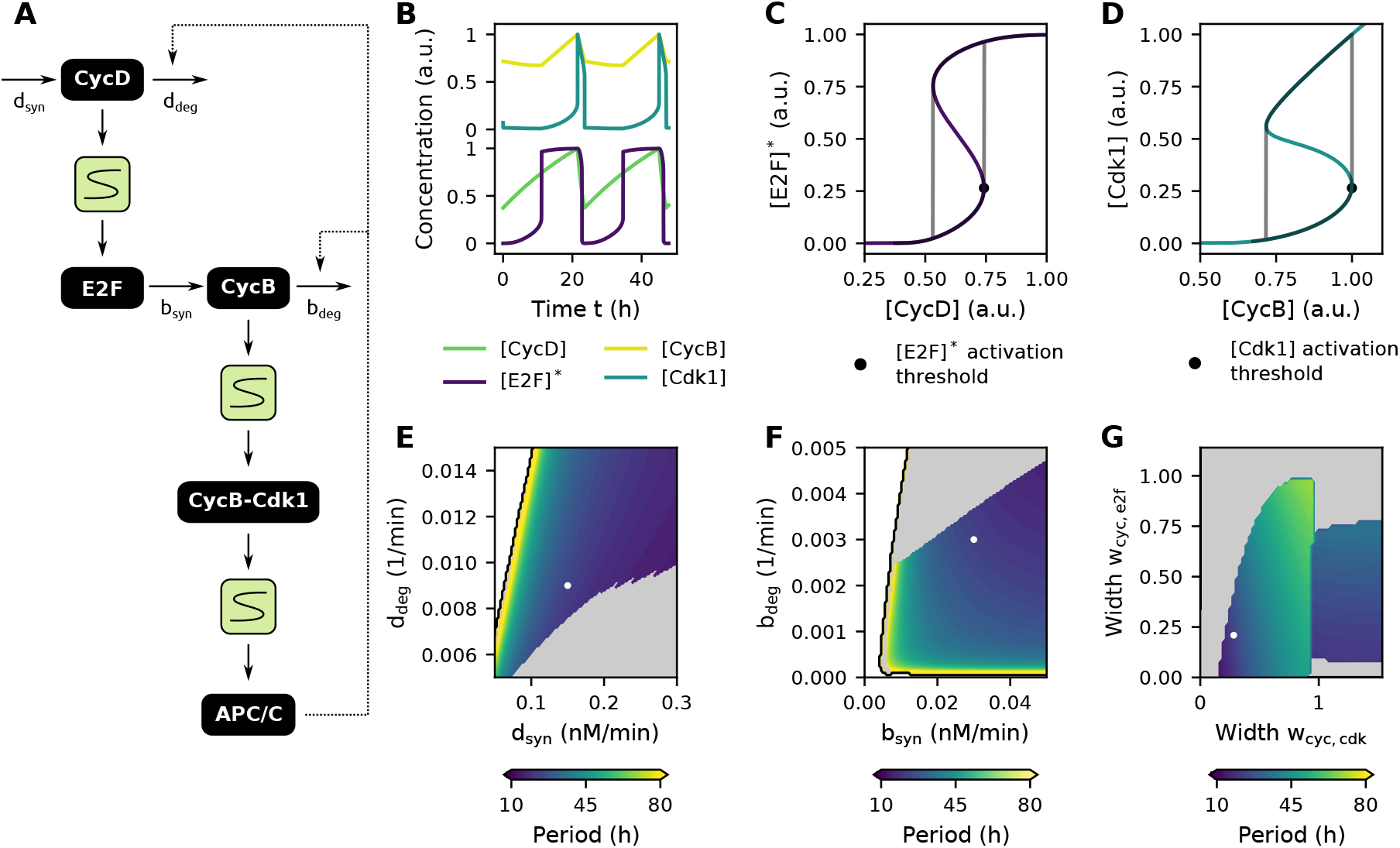
The cell cycle can be represented as a chain of interlinked bistable switches. Parameter values from Table 2 or as indicated in the figure. **(A)** Block scheme of interlinked S-shaped modules representing the overall cell cycle. **(B)** Oscillating time traces of the cell cycle model. **(C,D)** Time traces from B orbiting around the bistable response curves. As in B, concentrations are normalized for their maximal value. **(E,F,G)** Overall period of the oscillations as a function of CycD (E) or CycB (F) synthesis and degradation rates and the width of the S-shaped responses shown in C and D (G). No oscillations exist in the white regions. Grey regions represent oscillations that are not orbiting around all three bistable switches. White dots denote parameter values used in B-D.

As seen from Fig 8B, this modular representation of the cell cycle can produce oscillations in the concentration levels of the biochemical components. Synthesis of [CycD] (here used to directly model the effect of external growth factors), results in a linear increase of its concentration levels. Once [CycD] levels reach the [E2F]* activation threshold, [E2F]* activity suddenly rises (Fig 8B,C). Activated [E2F]* then (indirectly) induces the expression of [CycB], followed by the activation of [Cdk1] when [CycB] levels cross the [Cdk1] activation threshold (Fig 8B,D). As in the previous model of the G2-M transition, active CycB-Cdk1 induces APC/C activation and subsequent degradation of the cyclins (Fig. 8C,D, S1 Video).

**Table 2.** Default parameter values for modeling the cell cycle as interlinked bistable switches.

| Parameter | Value | Units | Description |
| --- | --- | --- | --- |
| $r$ | 0.5 | | As defined in $\xi$ |
| $\alpha_{e2f}$ | 5 | | As defined in $\xi$ |
| $\alpha_{cdk}$ | 5 | | As defined in $\xi$ |
| $\alpha_{apc}$ | 5 | | As defined in $\xi$ |
| $n$ | 15 | | Hill coefficient |
| $\epsilon_{e2f}$ | 0.01 | | Time constant E2F reaction |
| $\epsilon_{cdk}$ | 0.01 | | Time constant Cdk1 reaction |
| $\epsilon_{apc}$ | 0.01 | | Time constant APC/C reaction |
| $\delta_d$ | 0.05 | | Basal CycD degradation |
| $\delta_b$ | 0.05 | | Basal CycB degradation |
| $K_{cyc,e2f}$ | 120 | nM | Threshold of E2F activation by CycD |
| $K_{cyc,cdk}$ | 40 | nM | Threshold of Cdk1 activation by CycB |
| $K_{cdk,apc}$ | 20 | nM | Threshold of APC/C activation by Cdk1 |
| $d_{syn}$ | 0.15 | nM/min | Synthesis rate of CycD |
| $d_{deg}$ | 0.009 | 1/min | Apparent first order degradation of CycD |
| $b_{syn}$ | 0.03 | nM/min | Synthesis rate of CycB |
| $b_{deg}$ | 0.003 | 1/min | Apparent first order degradation of CycB |

The oscillatory behavior of this system depends on how the time traces orbit around the steady-state response curves, and hence are determined by the synthesis and degradation rates of the cyclins, together with the shape of the response curves themselves. For certain combinations of parameters, no oscillations can be observed (white regions in Fig 8E-G). For other combinations oscillations do exist, but these do not circle around all three response curves (grey regions in Fig 8E-G; the system might for example move along the top branch of the [E2F]* response curve, while orbiting around the [Cdk1] nullcline), resulting in irregular oscillatory patterns and hence irregular cell cycle progression (S7 Fig). The duration of the different phases depends on the values of production and degradation rates. A smaller synthesis rate *d*_syn_ of [CycD] and larger degradation rate *d*_deg_, entails a longer time needed for [CycD] levels to reach the threshold value for [E2F]* activation. This leads to oscillations with elongated G1 phases and thus longer overall periods (Fig 8E and S8 Fig). A decrease in the synthesis rate *b*_syn_ or degradation rate *b*_deg_ of [CycB] brings about oscillations with increased periods (Fig 8F). Whereas the effect of the synthesis rate can mainly be attributed to an elongation of the S-G2 phase, a diminished degradation rate causes an extension of the M phase (S8 Fig).

### Control mechanisms affecting the bistable switches shape the dynamics of the cell cycle

Unlike the early embryonic cell cycle, the somatic cell cycle is no ‘clock-like’ oscillator with a fixed period, but instead a ‘domino-like’ oscillator in which tight control mechanisms or ‘checkpoints’ have been established that safeguard correct DNA replication and cell division. Hence, the cell cycle representation from the previous section — in which no such checkpoints were considered — needs to be refined. Interestingly, the bistable nature of the cell cycle transitions itself can provide the means for this regulation by shifting the (in)activation thresholds (i.e. folding points) of the S-shaped response curves to lower or higher levels [13, 69, 70].

As a first example of a checkpoint, let us consider the so-called restriction point (RP). The traditional view of this checkpoint states that cells in G1 phase are predetermined to stay in this phase and not enter S phase. Only if sufficiently high concentrations of growth factors are present in the extracellular environment, leading to sufficiently high synthesis rates of CycD, the cell will progress into S-phase and irreversibly commit to complete the started round of cell division [71]. Interlinked bistable switches can account for such behavior as — once activated — [E2F]* activity can remain high even if [CycD] synthesis is strongly reduced (grey area in Fig 9A). This high [E2F]* activity then further induces [CycB] expression to reach the [Cdk1] activation threshold followed by activation of [APC]*.

**Fig 9.**
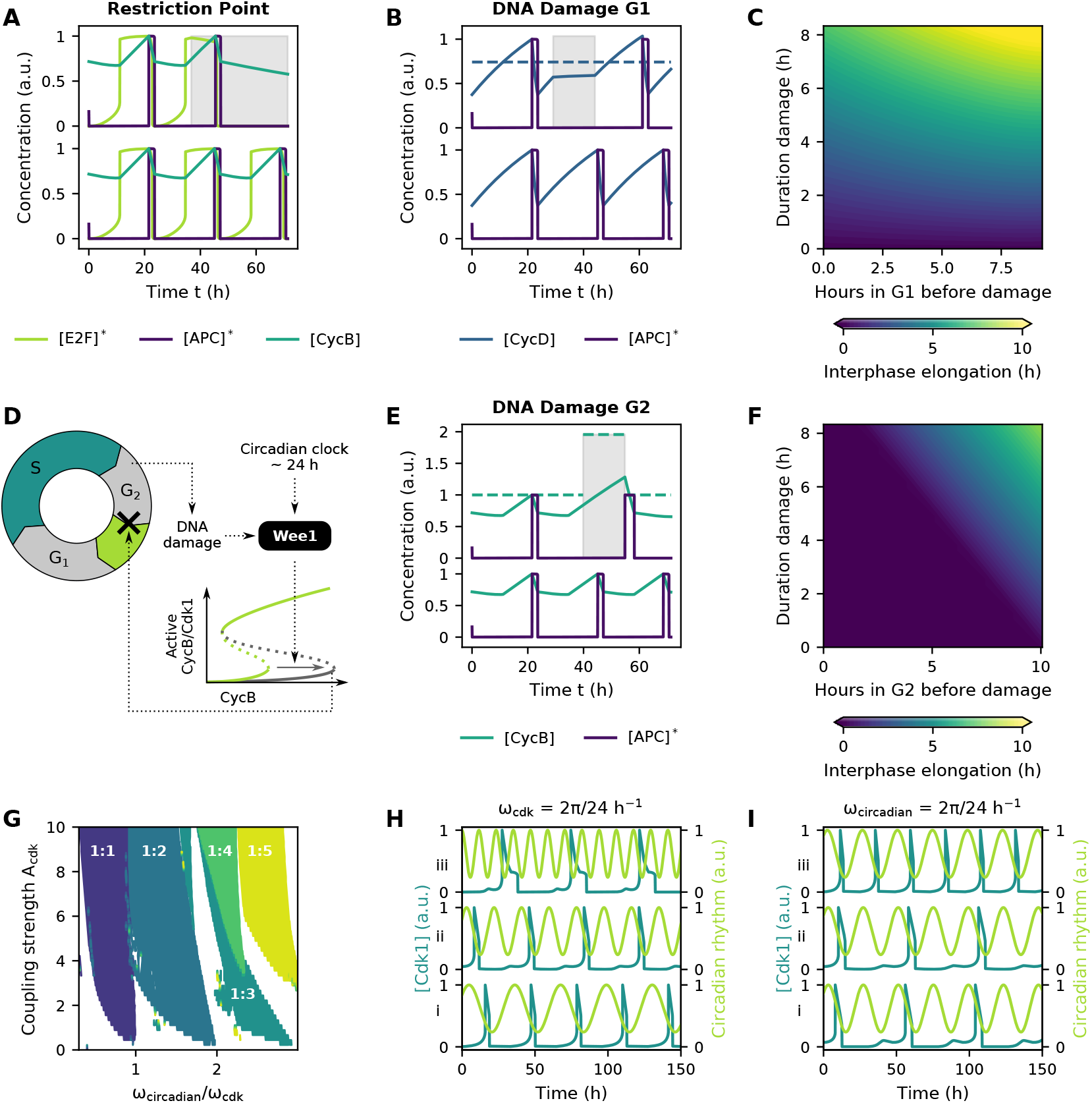
Control mechanisms affecting the bistable switches shape the dynamics of the cell cycle. Parameter values as in Table 2 or as indicated. **(A)** Time traces comparing unperturbed cell cycle progression (bottom) with progression after reduction of the [CycD] synthesis rate. In line with the classical interpretation of the RP, a started round of cell division can still be completed (as seen from the activation of [APC]*) even if [CycD] synthesis is reduced (grey area). **(B)** Time traces comparing unperturbed cell cycle progression (bottom) with progression after DNA damage in G1 phase. Dotted line represents the [E2F]* activation threshold and grey area marks region of increased [CycD] degradation. **(C)** Elongation of interphase after DNA damage in G1. **(D)** The kinase Wee1 is controlled by DNA damage and circadian rhythms and affects the Cdk1 activation threshold. **(E)** Time traces comparing unperturbed cell cycle progression (bottom) with progression after DNA damage in G2 phase. Dotted line represents the [Cdk1] activation threshold. Similar as in A and B, concentrations are normalized for their maximal value in the unperturbed case. **(F)** Elongation of interphase after DNA damage in G2. **(G)** Arnold tongues showing p:q phase locking between [Cdk1] oscillations and the circadian clock. The coupling strength *A*_cdk_ is a measure for the extent to which the [Cdk1] activation threshold is shifted to higher [CycB] levels. **(H)** Time traces of phase locking for a constant natural frequency of the cell cycle (i.e. 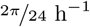) but different frequencies of the circadian clock ((i) 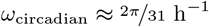, (ii) 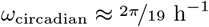, (iii) 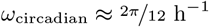. **(I)** Time traces of phase locking for constant circadian frequency (i.e. 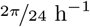) but different natural frequencies of the cell cycle ((i) 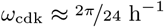, (ii) 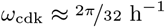, (iii) 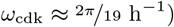). A coupling strength of *A*_cdk_ = 8 was used in H and I.

A second example of cellular control is the way cells preserve genomic integrity by arresting the cell cycle whenever deleterious DNA lesions are encountered. During G1 phase, DNA damage can prevent further cell cycle progression by inducing the degradation of CycD [72]. This behavior can be understood in our model by looking at the bistable switch governing the G1-S transition. The increased degradation of [CycD] (grey area in Fig 9B) keeps its levels below the [E2F]* activation threshold, thus blocking S phase entry, until DNA repair re-establishes normal degradation rates and [CycD] levels can rise beyond the threshold. The extent to which interphase is lengthened depends on when in G1 the damage was imposed, the duration of the damage and the time required to reach the [E2F]* activation threshold after the damage was repaired (Fig 9B,C).

DNA repair mechanisms during G2 phase have been associated with the activation of the kinase Wee1, which plays an important role in inactivating CycB-Cdk1 complexes and thus preventing entry into M-phase (Fig 9D) [73]. Exploiting the fact that increased Wee1 activity shifts the Cdk1 activation threshold to higher CycB levels [13, 59], the effect of DNA damage can straightforwardly be implemented by tuning the *α* parameter of the S-shaped module. As long as DNA damage is present, the [Cdk1] activation threshold is shifted to the right (grey area in Fig 9E) and [CycB] levels keep rising until a steady state is reached. When extensive damage is encountered, a permanent raise of the threshold might result in permanent cell cycle arrest. However, if DNA damage is repaired, Wee1 gets inhibited again, the activation threshold shifts back to lower [CycB] levels and [Cdk1] is activated, resulting in subsequent [APC]* activation. From the time traces (Fig 9E), it can be observed that cells enter mitosis with higher [CycB] levels (and thus also higher [Cdk1] levels) when recovering from DNA damage in comparison with unperturbed cells. Consequently, a longer time period is required to reach the [APC]* inactivation threshold and M-phase is prolonged. In contrast with the DNA checkpoint in G1 phase, modification of the bistable response curve in G2 can keep the total duration of interphase unaffected after DNA damage. If the damage is repaired before [CycB] levels reach the original [Cdk1] activation threshold, shifting it to higher levels has no effect on interphase duration (Fig 9F).

These examples show that a modular description can be used to implement biological events such as DNA damage in a phenomenological way: by adjusting the overall effect on the shape of the S-shaped response, without needing to know the exact molecular mechanisms. We provide another example of this approach by linking the cell cycle to the 24h circadian clock. Expression of the Wee1 kinase is not only affected by DNA damage but also tightly regulated by the Clock-Bmal1 transcription factor complex, a master regulator of the circadian clock. This couples the cell cycle to the circadian clock (Fig 9D) [74]. We explored the dynamics of such unidirectional coupling by introducing the circadian regulation of Wee1 into the model. More specifically, the [Cdk1] activation threshold — which is regulated by Wee1 — was periodically shifted between its basal [CycB] level and a predefined maximal level by periodically altering parameter *α*_cdk_ (see Methods). When two oscillators are coupled like this, one can expect to observe so-called p:q phase locking, meaning that p cycles of the cell cycle are completed for q cycles of the circadian clock. Whether such behavior indeed occurs depends on two factors [75]: (1) the relative frequencies of the circadian clock and the unforced cell cycle (i.e. the frequency in the absence of any circadian control, also called the natural frequency), and (2) on the strength of the coupling (here *A*_cdk_, i.e. the extent to which the [Cdk1] activation threshold is shifted to higher [CycB] levels over one period). When plotting regions of locking as function of the frequencies and coupling strength, one often finds so-called Arnold tongues (see Methods).

Typically, Arnold tongues span wedge-shaped regions in parameter space (at least below a certain critical coupling strength): for low coupling strengths, p:q phase locking would only occur when the natural frequency of the cell cycle is close to the frequency of the circadian clock, whereas a larger mismatch between the two frequencies may still result in synchronization if the coupling strength increases. In our case, we do find these regions, but the Arnold tongues solely widen to the left hand side (Fig. 9G). This indicates that the circadian clock only seems capable of lengthening the cell cycle and not shorten it. Indeed, examining the time traces in Fig. 9H, a 24h natural period of the cell cycle is elongated till 31h (i), 38h (ii) and 47h (iii) by a circadian rhythm of 31h, 19h and 12h respectively (see S9 Fig for additional time traces). Similarly, in Fig. 9I, a constant circadian rhythm of 24h elongates cell cycles with a natural period of 24h (i), 32h (ii) and 19h (iii) to forced periods of 48h, 48h and 24h, respectively. Considering that the circadian clock in our model only shifts the [Cdk1] activation threshold to higher [CycB] levels, relative to those of the unforced cell cycle, explains why the forced cell cycle period cannot become shorter than the unforced one. Of note, this observation is in agreement with the findings in [76], where unidirectional regulation of Wee1 by the circadian clock was analyzed using a mechanistic model. There too, Arnold tongues were found to only widen to the left, at least when a basal Wee1 synthesis rate was included.

## Discussion

In an attempt to unravel the complexity of living systems, it is convenient to envision them as a hierarchical structure of interlinked biological ‘modules’. Under the premise that the functionalities of these individual modules do not change when combined with each other, knowing their behavior allows to predict the behavior of the overall system [35]. Two proteins (or protein domains), for example, can be combined to generate recombinant proteins such as fused fluorescent reporters. Additionally, transcriptional promoters can be combined in synthetic networks that possess desired dynamical features, such as the oscillatory behavior of the ‘repressilator’, which consists of three negative feedback modules [77]. Although many examples of modularity exist, it has been recognized for a long time [78] that complex living systems can not always be understood as the sum of their constituent parts. Indeed, mechanisms have been described that can hamper the validity of a modular approach, such as off-target effects of transcription factors or the competition for cellular resources. At the same time however, methods to overcome these problems in engineered biological circuits have been proposed [35].

To gain insight into the dynamics of individual or interlinked modules that are difficult to understand in an intuitive way, mathematical models can be very useful. As such, they also provide helpful tools to guide the design of synthetic biological systems [79]. One way to set up a mathematical model is to start from known biochemical interactions and use the law of mass action to derive kinetic rate equations for the concentrations of the biochemical components. However, such mechanistic models often contain a large numbers of variables and parameters, many of which can be difficult to measure experimentally. This can impede generalization of the obtained results, which might strongly depend on the particular set of chosen parameter values. Therefore, techniques for reducing the complexity of mathematical models while preserving their most fundamental characteristics have been a topic of great interest for several decades, ranging from the well-established quasi-steady-state assumption to more advanced mathematical algorithms [80–83].

A conceptually easy approach to reduce the complexity of biochemical equations is to replace the detailed reaction mechanisms of certain biological modules by a mathematical function that explicitly describes their resulting dynamical behavior. Such a functional or phenomenological approach not only reduces the complexity of mathematical models, but it can also be a valuable alternative to model biochemical networks for which the underlying interaction patterns have not been identified completely. Furthermore, parameters of functional modules correspond directly to observable measurements that are often easier to determine experimentally than the values of biochemical rate constants. In our example of a bistable switch, the parameter *α* correlates with the width of the bistable region. By choosing the function *ξ* appropriately, the activation and inactivation threshold can thus be set directly in accordance with a response curve obtained in the lab. Moreover, the fact that these features are directly tunable allows us to answer questions such as ‘how does the activation threshold influence the period of oscillations?’. In a mechanistic model, answering such questions would depend on knowledge about what biochemical parameters determine the activation threshold, which might be less straightforward. The same argument holds for other properties, such as steepness of a response or a time delay. Thus, a direct intuitive link between the model and experimentally determined response curves exists when using functional modules. This is an important advantage of this strategy over other frameworks that are suitable for modeling biochemical reactions without considering mechanistic details, such as Boolean networks [84], especially for biologists without mathematical background.

Functional modules based on the Hill equation have been extensively used to describe steep sigmoidal response curves, which can originate from various biochemical mechanisms (e.g. multisite phosphorylations, cooperative binding events and stoichiometric inhibition). All of the molecular details that may generate such a response are then hidden in a few parameters such as the exponent *n* and a threshold *K*. In contrast, S-shaped response curves are typically not included explicitly in mathematical models. An important reason for this might be that an S-shaped response is not a function, because there are multiple output values for a single input. One contribution of this paper is to show that, in fact, a Hill function can be smoothly changed into an S-shaped response curve by making the threshold dependent on the output variable. In this way, the equation

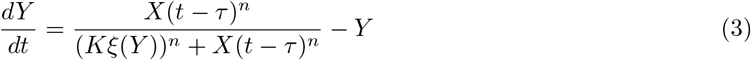

provides an easy way to model a system with input *X* and output *Y* that includes the three modules of ultrasensitive, S-shaped and time delayed responses. If the function *ξ* is equal to 1, we recover an ultrasensitive response for the steady state of *Y* as function of *X*. By choosing and tuning *ξ*, many different shapes of S-shaped response curves can be obtained. Whenever an experimentally determined steady state curve is available, the function *ξ* could also be fitted to such data (similar as what we did when fitting the S-shaped module to the mass-action model).

In the first part of the paper, we used this extended Hill function in combination with a negative feedback loop to model the early embryonic cell cycle oscillator and showed how the shape of the steady state response affects the oscillations. For ultrasensitive responses, no oscillations can be observed unless a sufficiently large time delay is present (S2 Fig, [60]). On the other hand, a system containing a S-shaped module readily sustains oscillations. Whether oscillations occur depends on the width of the S-region, and oscillations are more likely if there are time delays in the system. A second bistable switch can also facilitate oscillations, as we showed in a model that incorporates the two mitotic switches in the cell cycle. Furthermore, we demonstrated that the S-shaped module can faithfully reproduce the oscillatory dynamics of more detailed mass-action models.

In the second part, we extended the phenomenological model of the early embryonic cell cycle with an additional bistable switch for the G1-S transition. Furthermore, these bistable switches were exploited to model several cell cycle checkpoints, thus making the model representative for the somatic cell cycle. Even if the molecular underpinnings of these checkpoints are not incorporated into the model, the functional approach can give some basic understanding of events such as the restriction point (RP), DNA damage checkpoints and even coupling of the cell cycle with the circadian clock.

The restriction point as depicted here corresponds to the classical interpretation where newborn cells have to cross a threshold in order to activate E2F and commit to S-phase. It should be noted that such representation is a simplification, as it is known that additional bistable switches control entry into S-phase [85–87]. Furthermore, some cells retain hyperphosphorylated Rb proteins, which are incapable of inhibiting E2F, after division. These cells can directly start S-phase even in suboptimal conditions, challenging the classical view of the RP [71]. Interestingly, irregular cell cycle progression in which E2F activity remained high could also be observed for several combinations of parameters in our phenomenological model (S7 Fig). It would be interesting to see whether the biochemical mechanisms enabling cells to bypass the RP correspond with changes in functional responses identified here.

By implementing DNA damage in G2 via manipulation of the Cdk1 activation threshold, we found that cells enter M-phase with higher CycB-Cdk1 levels after DNA damage in G2 phase. This is in line with recently published experimental results [88]. Whether the subsequent M-phase takes longer than in unperturbed cells, as is predicted by the model, was not assessed in the cited study. If this would not be the case, it could indicate that either CycB degradation is accelerated after DNA damage or that the inactivation threshold too is shifted to higher CycB levels. Another interesting study which looked at the effects of DNA repair mechanisms was published by Chao *et al.*, [89]. There, the authors found that elongation of interphase after DNA damage in G2 depends on the severity of the damage, but is independent of when the cells were damaged within G2. This finding is in contrast with our model predictions that DNA damage in G2 can actually be repaired without interphase elongation. It should be noted that here we supposed that DNA damage can be detected and repaired at any time during G2, not only at a checkpoint at the end of G2 phase. Furthermore, our model does not account for regulatory processes such as decreased CycB synthesis [90] or increased CycB degradation [91], which might explain the discrepancies. Even if we cannot account for molecular details and nuances, the possibility to incorporate DNA damage in a phenomenological manner can be helpful to understand these events in a dynamical way.

Although in this work we combined the S-shaped/ultrasensitive/delay modules to generate a phenomenological model of the overall cell cycle, these modules are also applicable for modelling other processes based on their experimentally determined response curves. Bistable switches for example, are known to play a role during epithelial-to-mesenchymal transition [92] and differentiation in cell types ranging from embryonic stem cells [93] till osteogenic precursors [94] and *Drosophila* eyes [95]. Furthermore, the convenient way of combining several functional modules can be exploited to reconstitute interaction patterns with complex dynamical behavior, such as a combination of the ‘repressilator’ with bistable switches that has been observed during neural tube development [96].

The modeling strategy based on functional modules put forth in this work fits in the mindset of reducing the complexity of mathematical models and has the advantage of preserving an intuitive link with the experimentally measured response curves. Furthermore, the modular approach provides the flexibility for combining the modules in a multitude of different ways. As such, it can provide a toolbox for other researchers who want to generate phenomenological modular models, not only of the cell cycle but also of many other biological processes.

## Materials and methods

### Converting an ultrasensitive function into a S-shaped one

Starting from an (increasing) ultrasensitive response curve, one can obtain a S-shaped response by shifting the lower and upper part of the curve to the right and left respectively. For APC/C, the ultrasensitive response curve can be expressed as a Hill equation representing the fraction of activated APC/C to total APC/C molecules as a function of [Cdk1] (Fig. 10A):

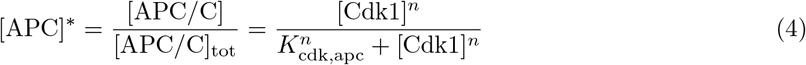

**Fig 10.**
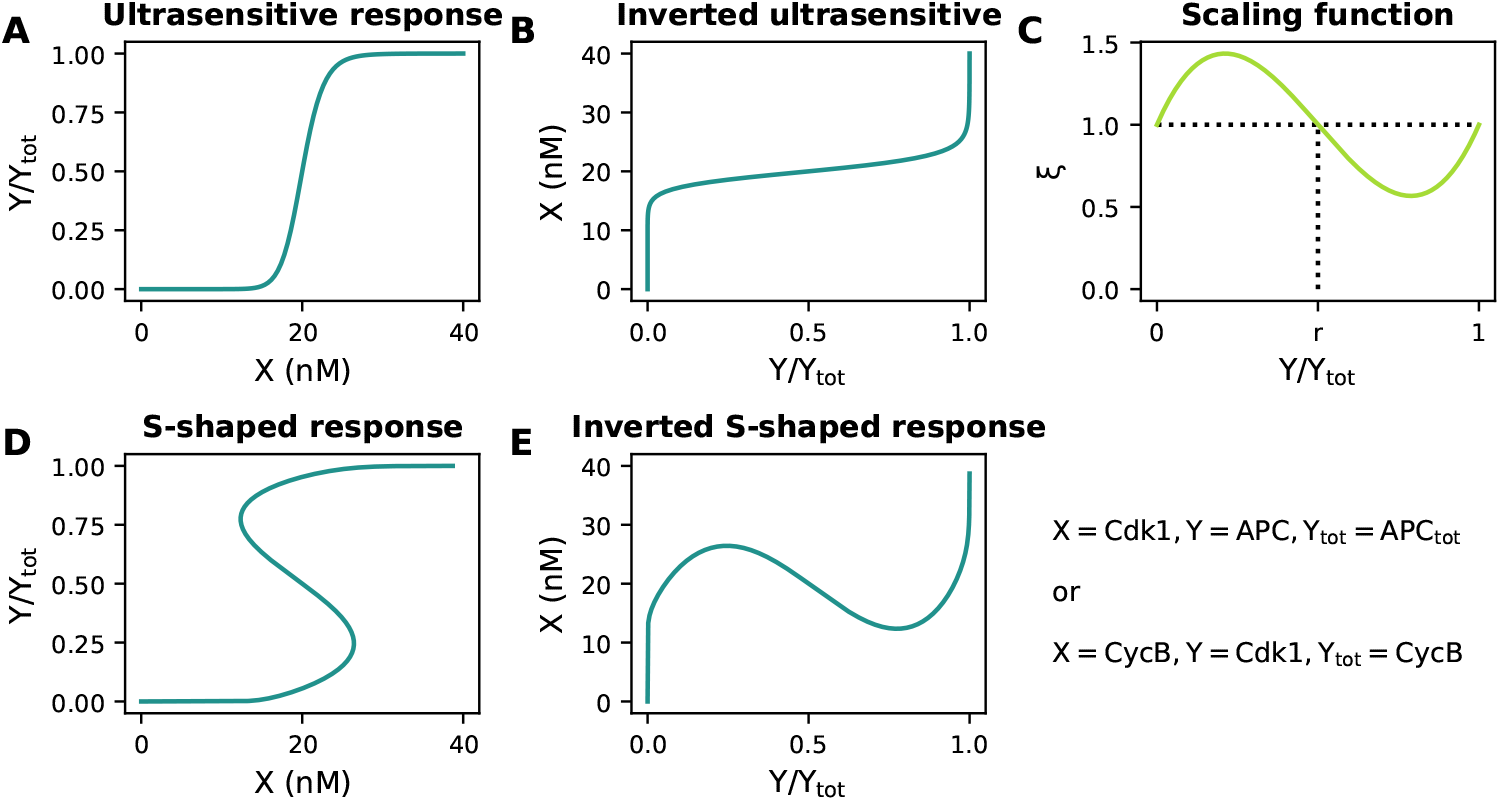
An ultrasensitive response curve (A) can be converted into a S-shaped one (D) by inverting the product (E) of a scaling function (C) with the inverted ultrasensitive response (B).

The S-shaped [APC]* response cannot be expressed as a function of [Cdk1] (since it would have multiple [APC]* values for a certain range of [Cdk1]). However, even if the function is *S*-shaped, there would be only one [Cdk1] value for each [APC]* value. This motivates us to invert the function (Fig. 10B):

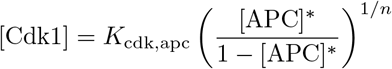

Subsequently, this inverted response can be multiplied by the scaling function *ξ*([APC]*) (Fig. 10C), such that [Cdk1] values increase for low [APC]* values and decrease for high [APC]* (Fig. 10E):

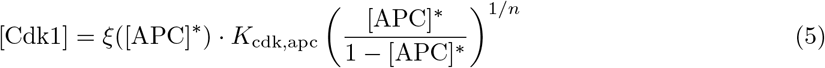

with *ξ*([APC]*) = 1+*α*_apc_·[APC]*([APC]* − 1)([APC]* − *r*). Rearranging Eq. 5 leads to an expression for a S-shaped response (Fig. 10D), analogous with Eq. 1 in the main text. This is not a function, but the expression can be put in a differential equation such that the steady state of this equation follows the S-shaped response:

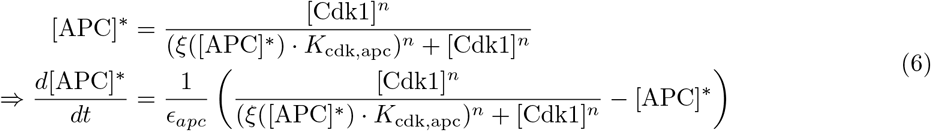

A similar approach can be followed for the bistable switch from total CycB-Cdk1 (i.e. [CycB]) to active CycB-Cdk1 (i.e. [Cdk1]). The ultrasensitive response is given by

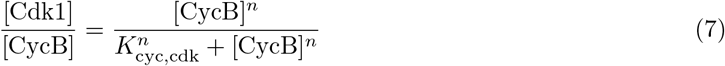

Rearranging and multiplying by 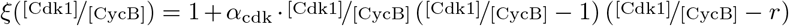 gives:

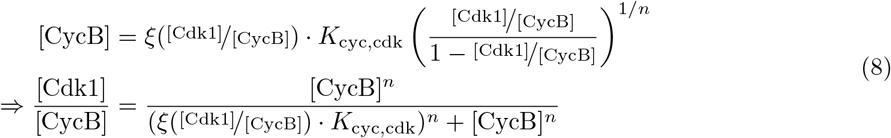

This means that

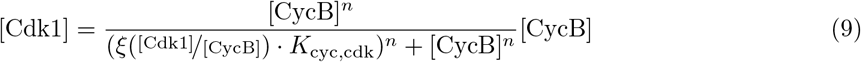

Again, this is not a function, but can be made the steady state of a differential equation (system (iii) in the main text):

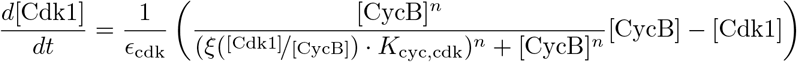

It needs to be emphasized that both Eq. 4 and 7 describe an ultrasensitive response of which the output is given by the concentration level of a compound *Y*, normalized for the total amount *Y*_tot_. Accordingly, both expressions are limited to the interval 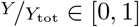. Similarly, Eq. 6 and 8 describe S-shaped responses for 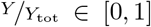. When incorporating these steady state expressions into the differential equations however, a distinction should be made. As [APC/C]_tot_ is a constant, the ratio 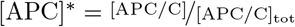 can directly be incorporated into the ODE for the rate of change of [APC]*. In contrast, [CycB] is not a constant and therefore, the rate of change of 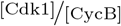 would need to be given by the quotient rule for differentiation, as both numerator and denominator are variables. Alternatively, [CycB] can first be moved to the right hand side of Eq. 8, resulting in Eq. 9. Subsequently, this expression for the steady state of [Cdk1] can be incorporated into an ODE describing the rate of change of [Cdk1]. Of note, Eq. 9 does no longer saturate at [Cdk1] = 1 (as was the case for Eq. 8 at 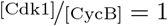) but keeps rising indefinitely (see for example Fig. 7C,E).

### Non-dimensionalization of system equations

To facilitate mathematical analysis, the system equations for the two-variable models described in the main text were non-dimensionalized by introducing the new variables:

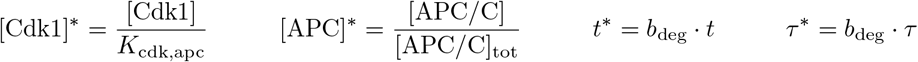

where *b*_deg_[min^−1^] is the reaction rate of the apparent first order degradation of CycB-Cdk1 complexes (i.e. variable [Cdk1]) by [APC]*. As an example, system (ii) then becomes:

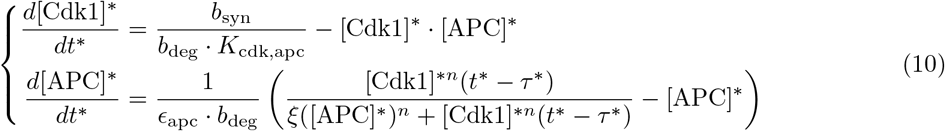

after which we can define 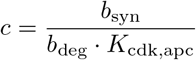, 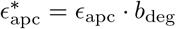 and *ξ*([APC]*) remains unaffected:

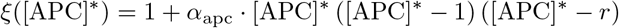

Standard parameter values used are: 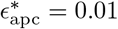, *n* = 15, *b*_deg_ = 0.1 min^−1^, *K*_cdk,apc_ = 20 nM, of which the latter three are based on experimental observations in *Xenopus laevis* eggs [12, 15, 30]. Obtained simulation results were scaled back to dimensional values where possible. For APC/C however, no experimental estimates for the total amount [APC/C]_tot_ were found and results are presented as [APC]*.

The system containing two S-shaped response (Eq. system (iii)) was non-dimensionalized in a similar way:

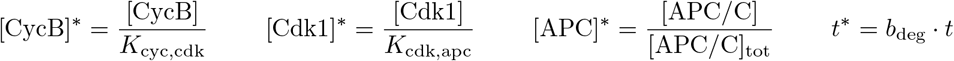

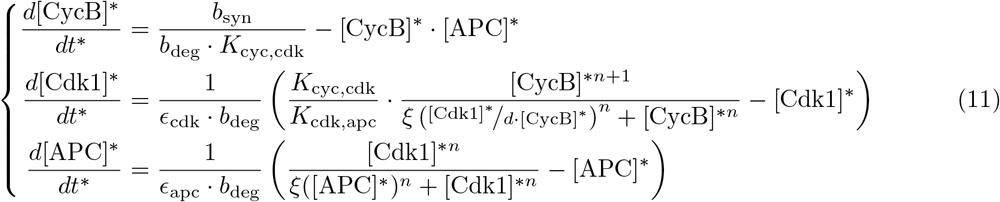

If 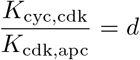, we get 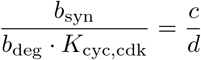. For the scaling functions we have *ξ*([APC]*) as before and

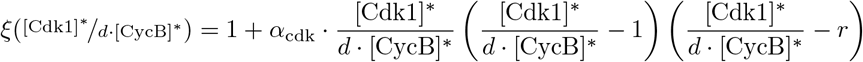

The same standard parameter values as the two-dimensional system were used, with the additional parameters being: *b*_deg_*ϵ*_cdk_ = *b*_deg_*ϵ*_apc_ = 0.01 and *K*_cyc,cdk_ = 40 nM [11].

### Conversion of parameter *α* to the width of the S-shaped region

All screens for which the width of the bistable region was altered, were performed by screening different values of *α* and linking this value to the width of the S-shaped region. The expression for the non-dimensionalized S-shaped response curve, i.e.

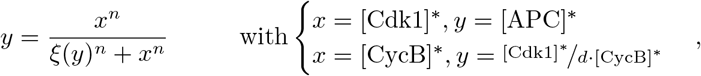

can be inverted, resulting in:

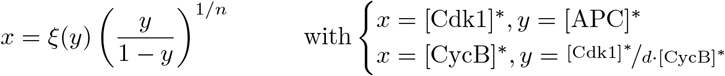

The width of the S-shaped region is given by the difference of *x*-values at the extrema of this inverted S-shaped response, which can be calculated as the roots of the derivative. For the cubic scaling function *ξ*(*y*) = 1 + *α*[*y*^3^ − (1 + *r*)*y*^2^ + *r* · *y*], we have:

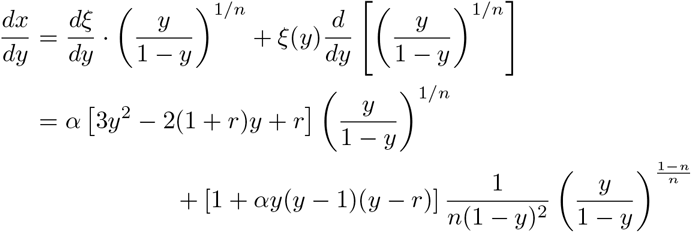

The roots of this function on the interval *y* ∈ [0, 1] were calculated numerically and used to determine the values of *x* at the extrema of the inverted S-shaped response. Either two extrema were found for which *x >* 0 (this was ensured by the choice of parameter *α*, see S1 Text), in which case the width of the S-shaped response curve was determined by their difference, or no extrema were found, in which case the width equaled zero.

### Mass-action model for the PP2A-ENSA-GWL network

The model of the PP2A-ENSA-GWL network was largely based on the network described in [61], where the double negative feedback between GWL and PP2A can give rise to bistability. GWL indirectly inhibits PP2A by phosphorylating ENSA, which is both a substrate and inhibitor of PP2A and binds it in a complex C [62]. Here, the model was converted into an oscillator by incorporating synthesis and degradation of [Cdk1]:

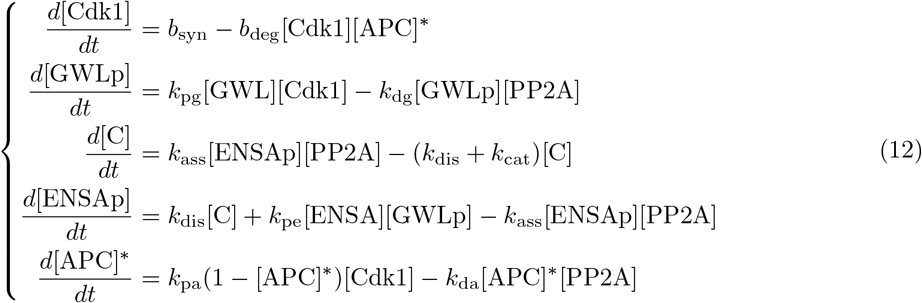

with conservation of mass giving:

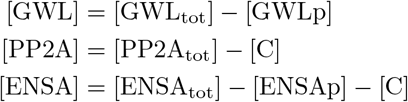

Parameter values were manually screened to obtain a steady-state response curve centered around [Cdk1] ≈ 20 nM (to be in line with *K*_cdk,apc_ = 20 nM used elsewhere in this paper) and obtain oscillations with biologically relevant periods (Table 1). Steady-state response curves were determined via a custom Python script performing numerical continuation [97].

### Fitting the S-shaped module to the mass-action model

Once the steady-state response of the mass-action model was determined, we manually fitted a piecewise linear scaling function *ξ*([APC]*) for use in system (i-b) (after non-dimensionalization).

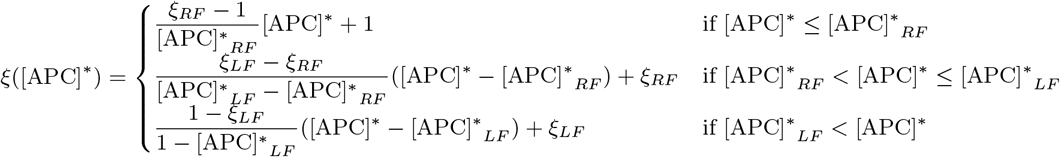

Here the subscripts RF and LF indicate right and left fold respectively (similar as in Fig. 4). For Fig. 5D, the best fit was obtained for [APC]*_*LF*_ = 0.65, [APC]*_*RF*_ = 0.39, *ξ*_*LF*_ = 0.76, and *ξ*_*RF*_ = 1.25. In Fig. 5G, the chosen values were [APC]*_*LF*_ = 0.39, [APC]*_*RF*_ = 0.16, *ξ*_*LF*_ = 0.73, and *ξ*_*RF*_ = 1.62. In each case, the location of the upper branch was manually shifted by multiplying the Hill-like term in system (i-b) (with *n* = 5) by a correction factor equaling 1 for Fig. 5D and 0.75 for Fig. 5G. For the time traces, 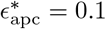 gave the best correspondence with the mass-action model.

### System equations for interlinked switches

The equations for the interlinked switches were derived following the same principles as described before, resulting in:

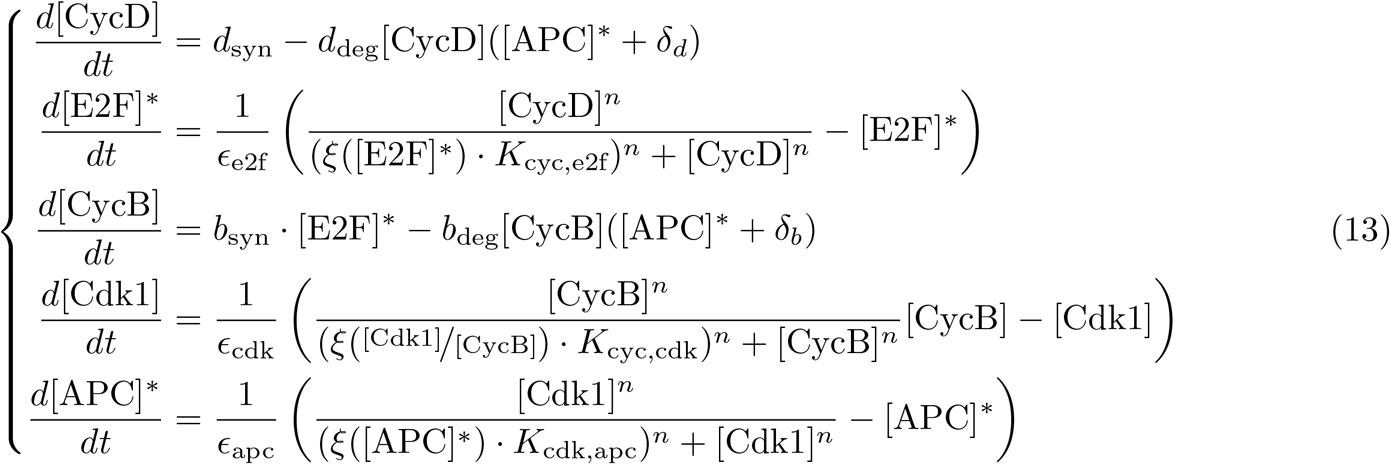

with

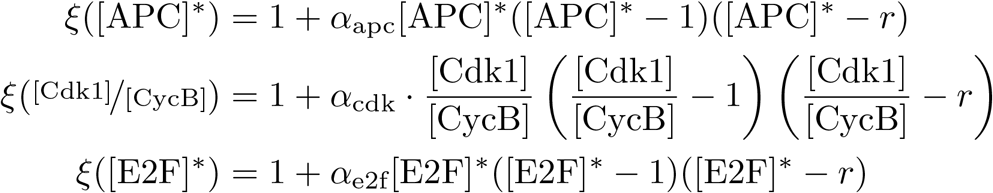

As before, 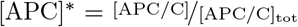 and similarly 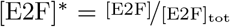. For all other variables and parameters, the original dimensions were retained. Default parameter values were manually screened so that oscillations with biologically relevant periods were obtained (Table 2). The threshold value *K*_cyc,e2f_ at which [CycD] activates [E2F]* was assumed to be 3 times higher than the threshold *K*_cyc,cdk_ at which [CycB] activates [Cdk1], i.e. 120 nM, based on simulations from [56] where CycD levels are about three times higher than CycB levels and on simulations from [19] where CycD levels rise up to ~ 100 nM.

To distinguish the different cell cycle phases in this model, cells were considered to be in M-phase wherever [APC]* > 0.95, in G1 wherever [E2F]* < 0.95 and [APC]* < 0.95, and in S/G2 wherever [E2F]* > 0.95 and [APC]* < 0.95.

To model the effect of the restriction point, reduced levels of external growth factors were incorporated into the model by reducing the synthesis rate *d*_syn_ of [CycD] by a factor 10. This number was chosen based on experimental measurements that demonstrated how growth factor stimulation increases CycD levels between 4 and 20 fold [98, 99]. DNA damage during G1 phase was accounted for by increasing the basal degradation rate of [CycD] (i.e. *δ*_*d*_) by a factor 3 and damage in G2 phase was modeled by increasing the width of the bistable [Cdk1] response (i.e. *α*_cdk_ = 30) to shift the right fold of the curve to higher [CycB] levels. The value of *α*_cdk_ = 30 was chosen so that the steady state of [CycB] in the standard model is below the [Cdk1] activation threshold.

### Simulations and analysis

All simulations were performed in Python 3.7. Ordinary differential equations were solved using the Python Scipy package solve ivp (method = Radau). For equations including a delay, the JITCDDE package was used [100]. The amplitudes and periods were numerically determined from the extrema in the time series via a custom Python script. Small amplitude oscillations and damped oscillations were omitted from the analysis.

### Arnold tongues and phase locking

Phase locking occurs when the ratio of periods (or frequencies) from two oscillators equals a rational number p:q, meaning that p cycles of one oscillator are completed while the second oscillator completes q cycles. To check for phase locking between the cell cycle and circadian clock, the system of interlinked switches (Eq. 13) was reanalyzed (again with parameters from Table 2), but this time the prefactor *α*_cdk_ in 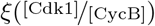 was periodically shifted between its basal level and a predefined maximal level (i.e. *α*_cdk_ + 2*A*_cdk_) at a forcing frequency *ω*_circadian_ ranging from 1/3 to 3 times the natural frequency of the [Cdk1] oscillations:

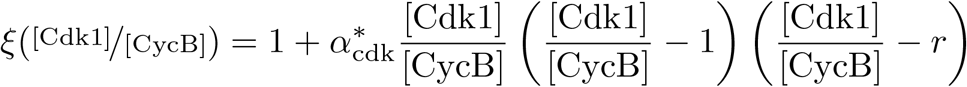

with 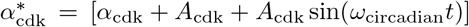. Custom Python code was used to find a repeating pattern in the forced [Cdk1] oscillations, based on the difference between the simulated time series and a time shifted version of itself (similar to calculating the autocorrelation). The ratio of the periods from this repeating pattern and the circadian clock was compared with p:q ratios for p and q in [1,2,3,4,5] to decide whether phase locking occurred.

## Supporting information

**S1 Text. Supplementary information.** This file contains additional mathematical analysis of the models and the supplemental figures listed below.

**S1 Fig Effect of parameters on the scaling function and system response.** Here we show in more detail how the different parameters affect the shape of the scaling function *ξ* and the overall dynamics of the system.

**S2 Fig Oscillations for a time delayed, ultrasensitive cell cycle model.** Ultrasensitivity by itself cannot induce oscillatory behavior of a two-dimensional system, i.e. either bistability needs to be incorporated or a sufficiently large time delay. In the main text, we focused on the effect of bistability, whereas here we summarize how ultrasensitivity in combination with a time delay can sustain oscillations.

**S3 Fig Oscillation period and amplitude for the bistable module.** The period and amplitudes of [Cdk1] and [APC]* oscillations for system (i-b) as a function of the relative synthesis *c* and the width of the bistable response curve. Related to Fig 3 in the main text.

**S4 Fig Alternative definitions of the scaling function *ξ***. Definitions for the scaling function *ξ* other than a cubic function can be used. The concept of ‘symmetric’ and ‘asymmetric’ bistable nullclines as used in the main text is visualised based on some alternative *ξ* functions.

**S5 Fig Oscillation period and amplitude for the delayed bistable module.** The period and amplitudes of [Cdk1] and [APC]* oscillations for system (ii) as a function of the width of the bistable response curve and the time delay. Related to Fig 6 in the main text.

**S6 Fig The width of the bistable modules in the three dimensional model affects the amplitude of the oscillations.** In Fig 7 in the main text we showed how the [Cdk1] amplitudes change by altering the threshold values *K* of the bistable switches. Here, we compare these results for different widths of the bistable response curves and show how the amplitude is proportional to the bistable width.

**S7 Fig Irregular cell cycle oscillations in a chain of bistable switches.** In Fig 8 in the main text we indicated grey regions in parameter space for which irregular oscillations were observed. Here, we show time traces of [E2F]* for such irregular patterns.

**S8 Fig Effect of synthesis and degradation rates of CycD and CycB on the duration of different cell cycle phases.** In Fig 8 in the main text we showed the effect of changing synthesis and degradation rates on the overall length of the cell cycle. Here, we separate the effects on the different cell cycle phases.

**S9 Fig Phase locking between the circadian clock and the cell cycle.** Time traces showing the absence or presence of p:q phase locking between the cell cycle and the circadian clock for several parameter combinations. Related to the Arnold tongues shown in Fig 9.

**S1 Video The cell cycle can be represented as a chain of interlinked bistable switches.** Video showing how the cell cycle progresses through the different bistable switches as shown in Fig 8.

## Acknowledgments

We are grateful to Prof. Catherine Verfaillie and the members of the Gelens lab for useful comments on the manuscript.

